# DNA barcode reference libraries for the monitoring of aquatic biota in Europe: Gap-analysis and recommendations for future work

**DOI:** 10.1101/576553

**Authors:** Hannah Weigand, Arne J. Beermann, Fedor Čiampor, Filipe O. Costa, Zoltán Csabai, Sofia Duarte, Matthias F. Geiger, Michał Grabowski, Frédéric Rimet, Björn Rulik, Malin Strand, Nikolaus Szucsich, Alexander M. Weigand, Endre Willassen, Sofia A. Wyler, Agnès Bouchez, Angel Borja, Zuzana Čiamporová-Zaťovičová, Sónia Ferreira, KD Dijkstra, Ursula Eisendle, Jörg Freyhof, Piotr Gadawski, Wolfram Graf, Arne Haegerbaeumer, Berry B. van der Hoorn, Bella Japoshvili, Lujza Keresztes, Emre Keskin, Florian Leese, Jan Macher, Tomasz Mamos, Guy Paz, Vladimir Pešić, Daniela Maric Pfannkuchen, Martin Andreas Pfannkuchen, Benjamin W. Price, Buki Rinkevich, Marcos A. L. Teixeira, Gábor Várbíró, Torbjørn Ekrem

## Abstract

Effective identification of species using short DNA fragments (DNA barcoding and DNA metabarcoding) requires reliable sequence reference libraries of known taxa. Both taxonomically comprehensive coverage and content quality are important for sufficient accuracy. For aquatic ecosystems in Europe, reliable barcode reference libraries are particularly important if molecular identification tools are to be implemented in biomonitoring and reports in the context of the EU Water Framework Directive (WFD) and the Marine Strategy Framework Directive (MSFD). We analysed gaps in the two most important reference databases, Barcode of Life Data Systems (BOLD) and NCBI GenBank, with a focus on the taxa most frequently used in WFD and MSFD. Our analyses show that coverage varies strongly among taxonomic groups, and among geographic regions. In general, groups that were actively targeted in barcode projects (e.g. fish, true bugs, caddisflies and vascular plants) are well represented in the barcode libraries, while others have fewer records (e.g. marine molluscs, ascidians, and freshwater diatoms). We also found that species monitored in several countries often are represented by barcodes in reference libraries, while species monitored in a single country frequently lack sequence records. A large proportion of species (up to 50%) in several taxonomic groups are only represented by private data in BOLD. Our results have implications for the future strategy to fill existing gaps in barcode libraries, especially if DNA metabarcoding is to be used in the monitoring of European aquatic biota under the WFD and MSFD. For example, missing species relevant to monitoring in multiple countries should be prioritized. We also discuss why a strategy for quality control and quality assurance of barcode reference libraries is needed and recommend future steps to ensure full utilization of metabarcoding in aquatic biomonitoring.

## 1. Introduction

### 1.1 DNA barcoding for monitoring aquatic life

Aquatic life is of central importance to human well-being and essential for our understanding of natural history, evolution and ecology. From the deepest oceans to the highest peaks, life in water characterizes environmental conditions, and constitutes invaluable ecosystem functions with services for a wide array of communities (Borgwardt et al., 2019; Rouillard et al., 2018). For these reasons, our ability to assess aquatic biodiversity and monitor its change over time is of great significance, not only to prevent biodiversity loss, but to ensure our own welfare.

The world’s oceans cover 70% of the Earth’s surface and are home to approximately 242,000 described species (Horton et al., 2018). It is estimated, however, that 91% of eukaryotic marine life is undescribed, and that the total number of marine species is around 2.2 million (Mora et al., 2011). More than one third of the world’s human population lives in the coastal zone, and ecosystem services provided by the marine environment are both crucial to human well-being and affected by our activities (Barbier, 2012; Barbier, 2017). In Europe, the Marine Strategy Framework Directive (MSFD, Directive 2008/56/EC) aims to achieve “good environmental status” of marine waters by 2020 and to protect marine environments in the European Union (European Commission, 2008). The MSFD includes a wide array of requirements in its ecosystem-based approach for assessment and monitoring, including information on animal and plant communities (Borja et al., 2013). A large percentage of undescribed biota certainly hampers community comparisons among sites and regions, and likely restrains the explanatory power of marine water quality indices (Aylagas et al., 2014).

Although representing only 0.01% of the Earth’s water, freshwater ecosystems hold about 6% of all described species (Balian et al., 2008; Dudgeon et al., 2006; Reid et al., 2018). Freshwater represents a valuable and irreplaceable natural resource, and scarcity as well as quality are likely to continue to affect the stability of human communities (Kreamer, 2012). Four-fifths of the world’s population now lives in areas where there is a threat to water security (UN World Water Assessment Programme, 2018), and it is estimated that demand for freshwater will increase by 20-30% by 2050 (Burek et al., 2016). Water quality as well as access to water is of global concern, and nature-based solutions have received increased attention as ways of improving water quality (UN World Water Assessment Programme, 2018). In Europe, assessments of water quality have been a hot topic for decades (Birk et al., 2012b; Hering et al., 2010; Leese et al., 2018; Metcalfe, 1989), and the use of biodiversity estimates for this purpose is central in the EU Water Framework Directive (WFD) (European Commission, 2000). Moreover, the highest proportion of species extinctions to date has been recorded in freshwater (Young et al., 2016), highlighting the importance of monitoring and protecting these ecosystems.

Thus, together with the Groundwater Directive (GWD, Directive 2006/118/EC) and the Habitats Directive (Directive 92/43/EEC), the WFD and MSFD make water quality monitoring of Europe’s aquatic environments legally binding in all EU member states, Norway, Iceland and Switzerland.

However, among countries there are large differences in the way biodiversity data are used to assess aquatic ecosystem quality status (Birk et al., 2012a; Kelly et al., 2015): different indices, different taxonomic groups, and different taxonomic levels are applied. Despite differences in methodology, the goals are similar and focus on the quantification of environmental states in comparison with reference conditions. Protocols and assessment metrics applied have undergone a sophisticated intercalibration procedure to harmonise data among countries and make ecological status assessments comparable.

To assess the ecological status, identification of aquatic organisms to family, genus or species-level by morphology is necessary, but it is not a straightforward process. For instance, individual differences in expertise, experience and opinion of the identifiers can result in different taxonomic groups being documented from the same waterbody, potentially leading to contrasting ecological assessments (Carstensen and Lindegarth, 2016; Clarke, 2013). An extensive audit of 414 macroinvertebrate samples taken as part of the monitoring programs of German rivers and streams (Haase et al., 2010) documented that 29% of the specimens had been overlooked by the primary analyst in the sorting stage, and that the identification of >30% of the taxa differed between the primary analyst and the auditors. Importantly, these results lead to divergent ecological assessments in 16% of the samples (Haase et al., 2010). Similar studies have been performed in Norway and Finland (Meissner et al., 2012; Meissner et al., 2017; Petrin et al., 2016) with comparable results. Despite the general challenges in using short, standardized molecular markers for identification (Hebert et al., 2016), DNA barcoding and metabarcoding offer a less subjective approach than morphology for the identification of organisms in aquatic assessments (Leese et al., 2018). Some issues still need to be solved and standard protocols to be developed before DNA metabarcoding becomes the method of choice in aquatic biomonitoring. The use of both organismal and environmental DNA (eDNA) in nature management decisions is already being tested in some European countries (Hering et al., 2018), and the genetic water quality index recently developed for marine waters (gAMBI) is performing well (Aylagas et al., 2018; Aylagas et al., 2014). The EU COST Action DNAqua-Net (CA15219) was initiated with the purpose of developing genetic tools for bioassessments of aquatic ecosystems in Europe (Leese et al., 2016). The network aims to evaluate existing methods and reference libraries, as well as to develop protocols and good practices in the use of DNA-based monitoring and assessments of aquatic habitats. By connecting scientists and stakeholders, DNAqua-Net so far has been a successful platform for this purpose.

Comprehensive DNA barcode reference libraries, such as the Barcode of Life Data Systems (BOLD (Ratnasingham and Hebert, 2007)) or GenBank (Benson et al., 2013), are essential for biodiversity monitoring if one wishes to utilise species’ autecological and biogeographic information gathered during the last century and to compare results with previous assessments. But also smaller, more taxon specific reference libraries, such as Diat.barcode library, formerly called R-Syst::diatom database (Rimet et al., 2016) are important as these might be easier to curate. Particularly in the current ‘big biodiversity data’ era, in which hundreds of millions of sequences can be generated during a single high-throughput sequencing run, we are no longer able to individually check sequence by sequence. It is thus imperative that effective quality filtering processes are embedded, including that reference libraries hold high standards and are well populated in order to trust (semi-)automated taxonomic assignments (Brodin et al., 2012; Carew et al., 2017; Ekrem et al., 2007; Hebert et al., 2003a; Mioduchowska et al., 2018; Porter and Hajibabaei, 2018). An elaborated quality assurance / quality control (QA/QC) system can serve both purposes. Building barcode libraries and associated voucher collections have therefore been major goals in individual projects as well as national barcode campaigns over the last decade. In Europe, some nations have been successful in obtaining funding to coordinate this work on a national level. Others have contributed to reference libraries on a project-by-project basis. The way the work on reference libraries has been organized is different between nations, and in some cases decisive for which taxonomic groups and regions were covered. We therefore find it informative and useful to briefly recapture the most important aspects of these initiatives in Europe.

### 1.2 Barcode Campaigns in Europe

The Austrian Barcode of Life (<u>ABOL</u>) is an initiative with the main aim to generate and provide DNA barcodes for all species of animals, plants and fungi recorded from Austria. The main purpose of the pilot phase (2014-2017) was to build up a network of biodiversity experts and conduct four pilot studies. Currently DNA barcodes are generated in a number of independently funded projects. The pilot phase and the continued coordination of ABOL is funded by the Ministry of Education, Science and Research and lies at the Natural History Museum Vienna. Apart from building up the reference library, necessary for genetically determining organisms, ABOL aims to stimulate biodiversity research by acquiring funds, fostering diverse applications of DNA barcoding, building up and exchanging skills within the network, and increasing public awareness for biodiversity.

The Finnish Barcode of Life (<u>FinBOL</u>) is a national project and a network of species experts with the goal of creating DNA barcodes for all species of animals, plants and fungi occurring in Finland. FinBOL has acted as a national node in the International Barcode of Life (iBOL) project. FinBOL has been funded almost continuously from 2011 by several national funding agencies. At the moment, FinBOL acts within the framework of the Finnish Biodiversity Information Facility (FinBIF) and is coordinated by the University of Oulu. DNA barcoding details for all Finnish species are provided in the <u>Laji.fi</u> portal, where progress is continuously updated. At present, over 100,000 specimens stored in Finnish collections have been subjected for barcoding, and DNA barcodes are available for about 20,000 species (~50%) reported from Finland. In the near future, FinBOL aims at broadening the nationwide DNA barcode reference library by adopting efficient high-throughput sequencing tools to recover sequence information from older museum specimens.

Since November 2011, the German Federal Ministry of Education and Research (BMBF) is funding a consortium of natural history museums and research institutions to set up the ‘German Barcode of Life’ initiative (<u>GBOL</u>). The main aim was to establish a network of professionals and non-professionals to start with the construction of a DNA barcode reference library for the fauna, flora and fungi of Germany. After the first phase (2011-2015) a national web portal for DNA barcodes and specimen data was developed and is continuously improved. It serves mainly the coordination of the collecting activities of over 250 scientists (amateurs and professionals) who provide their taxonomic expertise. In addition, more than 50 institution-based taxonomists contribute to GBOL. Of the 48,000 animal and 10,000 plant species (excluding algae and fungi) present in Germany, over 23,000 different species have been processed and DNA barcodes for them generated. In total, 295,000 specimens were submitted to GBOL institutes, and after choosing up to 10 individuals per species from throughout their distribution range in Germany, over 145,000 of them delivered a DNA barcode. The second phase of GBOL (2016-2019) has focussed on applications of DNA barcoding with dedicated PhD students working on specific aspects from metabarcoding for water quality assessments to developing a diagnostic microarray chip for the detection of phytopathogenic fungi. As a prerequisite for the successful implementation of the new techniques a core team and network of taxonomists is further expanding the reference library with DNA barcodes for another 13,800 species. With this target the database will be filled with about half of the known metazoan species of German animals and plants and be operable to identify the vast majority in terrestrial and aquatic environmental samples. Substantial contributions to the reference library for German taxa came from the project ‘Barcoding Fauna Bavarica (BFB)’, which started in 2009 and is supported by grants from the Bavarian State Government. The project focuses on animal biodiversity in Southern Germany and is coordinated by the Bavarian State Collection of Zoology (ZSM). Research activities involve close cooperation with the Biodiversity Institute of Ontario, which performs the sequence analyses under the framework of the International Barcode of Life Project (<u>iBOL</u>).

The Norwegian Barcode of Life Network (<u>NorBOL</u>) started in 2008 as a consortium of biodiversity institutions in formal agreement of advancing DNA barcoding in Norway. The four university museums in Bergen, Oslo, Tromsø and Trondheim have been hubs in the network since then, and together with the Biodiversity Institute of Ontario, Canada, the main partners in a national research infrastructure project that received funding from the Research Council of Norway and the Norwegian Biodiversity Information Centre (NBIC) in 2014. The major goal of the NorBOL-project was to database DNA barcodes of 20,000 Norwegian, Scandinavian or Polar species in BOLD by the end of 2018. However, also knowledge transfer, building expertise, and curation of specimen reference collections have been important tasks of the network. Close collaboration with the Norwegian Taxonomy Initiative, run by NBIC, has been crucial in this process as it has provided identified specimens of many organism groups available for DNA analysis. Several applied research and management projects have originated through collaboration in NorBOL.

The Swiss Barcode of Life (<u>SwissBOL</u>) is the national initiative for the creation of a genetic catalogue for all species occurring in Switzerland. SwissBOL officially started in 2012 supported by the Federal Office for the Environment, with the goal of establishing a network of scientists and institutions involved in the genetic inventory of Swiss biodiversity. During the pilot phase (2012-2015), 24 targeted projects were developed on different taxonomic groups: animals, plants, fungi, lichens and microorganisms. Ever since (transitory phase; 2016-2018), the coordination of SwissBOL has been funded almost continuously, and data has been acquired within only a few independently funded projects. In order to elaborate a national strategy for the development of projects generating novel genetic data, a non-profit association of experts was founded. Most recently, SwissBOL has been mostly working in the development of the concepts for the genetic database with the major goal of ensuring that the information related to the genetic data are accessible and linked together. The close collaboration with the GBIF Swiss Node (http://www.gbif.ch) has been fundamental to ensure the coherence of all the information provided with the standards defined at the national and international levels.

The Netherlands started their barcoding initiative NBOL for plants and animals in 2008, led by Naturalis Biodiversity Center in collaboration with a large number of Dutch NGOs and over 50 amateur naturalists. A considerable starting grant from the national government in 2010 gave a tremendous boost to the DNA barcoding infrastructure at Naturalis and hence to the national barcoding activities. So far, over 80,000 DNA barcodes have been generated. More than half of the barcodes have been uploaded to BOLD. However, most of these barcodes are still private because they are part of active research projects. Current barcoding efforts focus on the completion of reference libraries of freshwater and marine species (North Sea) for DNA-based biodiversity assessments, and are financed by private funding organisations.

Among various DNA barcoding initiatives in Portugal, one of the most prominent contributions has been provided by the network for barcoding marine life. This network was activated in 2008 through a research grant (LusoMarBoL-Lusitanian Marine Barcode of Life) from the national science funding body (Fundação para a Ciência e a Tecnologia - FCT), and has been active ever since through subsequent research grants. Core reference libraries for Portuguese marine life have been created, published and made available in BOLD, with particular focus on marine fish (Costa et al., 2012; Oliveira et al., 2016), annelids (Lobo et al., 2016), crustaceans (e.g. (Lobo et al., 2017) and molluscs (Borges et al., 2016).

While national DNA barcode initiatives often start opportunistically and register any species available for sampling, focus shifts to fill the gaps of the databases as soon as a critical number of species is registered. Which taxonomic groups have priority is typically connected to funded projects, available taxonomic expertise and scientific collections, and is not necessarily the same in each campaign. Among aquatic taxa, species-rich groups such as arthropods and polychaetes, or economically important groups such as fish, have seen some priority. However, when building barcode reference libraries, there has usually not been a general focus on species or organisms that are particularly relevant for water quality assessments towards WFD or MSFD from the start.

In addition to large national barcoding campaigns, smaller activities intended to generate reference barcodes of selected taxonomic groups (e.g. Trichoptera Barcode of Life), or regional biota (e.g. “Barcoding Aquatic Biota of Slovakia - AquaBOL.sk” and “Israel marine barcoding database”) exist. These initiatives, even if lacking substantial funding, can provide important data and in many cases be better targeted towards filling the gaps of barcode libraries than more general campaigns.

### 1.3 Biological Quality Elements

Different organism groups are used as Biological Quality Elements (BQEs) to assess the Ecological Quality Status (EQS) of aquatic ecosystems under the WFD. In the MSFD, biodiversity data in general, along with other related descriptors, are used to define Environmental Status (Borja et al., 2013; Zampoukas et al., 2014).

The MSFD is the first EU legislative instrument related to the protection of marine biodiversity. The directive lists four European marine regions: 1) the Baltic Sea, 2) the North-east Atlantic Ocean, 3) the Mediterranean Sea, and 4) the Black Sea. Member States of one marine region and with neighbouring countries sharing the same marine waters, collaborate in four Regional Sea Conventions (OSPAR^1^, HELCOM^2^, UNEP-MAP^3^ and the Bucharest Convention^4^). These different regions naturally share, or aim to share, taxa/species lists for biodiversity assessments and reporting status. The status is defined by eleven descriptors in the MSFD (e.g. biological diversity, non-indigenous species, fishing, eutrophication, seafloor integrity, etc.). For some descriptors, species ID is critical. National marine environmental monitoring often focuses on regular sampling sites and observations of specific habitats and its inhabitants, i.e. groups of organisms such as benthic macroinvertebrates, phytoplankton, or fish. As already mentioned, there exist large differences between countries in how biodiversity data are used to evaluate the quality status of aquatic ecosystems. This is indeed true for the marine environment, and only few countries were able to support this study with national taxalists directly associated to the MSFD. MSFD overlaps with WFD, and in coastal waters MSFD is intended to apply to the aspects of *Good Environmental Status* that are not covered by WFD (e.g. noise, litter, other aspects of biodiversity) (European Commission, 2017). In order to perform barcode gap-analyses for taxa of relevance to the directives and with a European marine perspective, we identified the possibilities of two existing taxalists: AZTI’s Marine Biotic Index (AMBI; (Borja et al., 2000)) and the European Register of Marine Species (<u>ERMS</u>).

The AMBI is used as a component of the benthic invertebrates’ assessment by several Member States in the four regional seas (Borja et al., 2009; European Commission, 2018), in the context of describing the sensitivity of macrobenthic species to both anthropogenic and natural pressures (see e.g. (Borja et al., 2000)). The index uses the abundance weighted average disturbance sensitivity of macroinvertebrate species in a sample (Borja et al., 2000), each species being assigned to one of five ecological groups (EG I-V; (Grall and Glémarec, 1997). The AMBI list includes approximately 8,000 taxa (only macroinvertebrates) from all seas, with representatives of the most important soft-bottom communities present at estuarine and coastal systems, from the North Sea to the Mediterranean, North and South America, Asia, etc. The second list used for the work is ERMS (Costello, 2000). This is a taxonomic list of species occurring in the European marine environment, which includes the continental shelf seas of Europe as well as the Mediterranean shelf, Baltic Seas and deep-sea areas (http://www.marbef.org/data/ermsmap.php) up to the shoreline or splash zone above the high tide mark and down to 0.5 psu salinity in estuaries. The register was founded in 1998 by a grant from the EU’s Marine Science and Technology Programme and contains tens of thousands of marine species, so for this study we used a relevant selection of organism groups within the register (see methods). In contrast to freshwater microphytobenthos, where ecological indices are calculated on the base of country specific index values attached to species names, marine microphytobenthos is not used for the calculation of ecological indices. And while all four regional sea conventions recognize the importance of marine microphytoplankton monitoring, no ecological index based on species-specific values is implemented. Monitoring of marine microphytoplankton is therefore carried out by monitoring the presence or abundance of all observable species as a biodiversity measure with an additional focus on the search for invasive species. This approach effectively extends the range of species monitored to the range of all known microphytoplankton species as there is no restriction to a list of species with ecological index values.

In freshwater, diatoms, with their huge species diversity, are particularly interesting ecological indicators (Stevenson, 2014). They have been routinely used for monitoring of surface waters for several decades (Rimet, 2012), and are required BQEs in assessments of surface waters in Europe and the United States (Barbour and United States. Environmental Protection Agency. Office of Water., 1999; European Commission, 2000). Until recently, the standardized methodology for biological monitoring using diatoms was uniquely based on microscopic determinations and counts (European Standard EN 14407:2014). This is quite time-consuming and requires expertise in diatom taxonomy; skills that can only be acquired after several months or years of practice. The development of high-throughput sequencing (HTS) technologies and DNA barcoding provides an alternative to the tedious work of morphological identification. The first proofs of concept, carried out on a few tens of samples, showed interesting and encouraging results (Kermarrec et al., 2013; Zimmermann et al., 2015). Recent studies confirmed that diatom indices obtained from DNA metabarcoding provide very similar results to diatom indices calculated by microscopic counts, both on a regional and national scale (Keck et al., 2017; Lefrancois et al., 2018; Rimet et al., 2018b; Rivera et al., 2018a; Rivera et al., 2018b; Vasselon et al., 2018; Vasselon et al., 2017). However, all these studies underlined the necessity of well-curated reference libraries. In Europe, efforts to develop such a resource are made by a group of diatom experts, which curate the Diat.barcode library (Rimet et al., 2016). They also proposed innovative methodologies based on HTS to fill the gaps of this database (Rimet et al., 2018a).

Aquatic macrophytes are recognized as a valid taxonomic group for assessing water quality according to the WFD. They reflect the morphological conditions of the water bodies (diversity and dynamics of the substratum, degree of rigid management of the banks) and are particularly interesting to assess nutrient pressure. Moreover, they react to anthropogenic interventions in the hydrological regime (potamalisation and water retention). Being plant organisms, macrophytes also present properties, such as longevity and immobility, that make them bad bioindicators in the short-term: they are able to integrate disturbed conditions over a considerably long period of time; it is impossible to accurately locate the source of pressures and the area of impact (Pall and Mayerhofer, 2015). According to the traditional definition, macrophytes are aquatic plants whose vegetative structure develops either in the water on a permanent basis or at least for a few months, or on the surface of water (Cook et al., 1974). These include species of the Charophyta (charales), the Bryophyta (mosses), the Pteridophyta (ferns) and the Spermatophyta (seed plants). In the present study we decided to focus our analyses on vascular plants only, which therefore regroups species from the divisions Pteridophyta and Spermatophyta. Concerning the choice of markers, DNA barcoding in plants is not as straightforward as in animals. The Consortium for the Barcode of Life (CBOL) Plant Working Group ended up by recommending the combination of two plastid loci for the standard plant barcode — rbcL+ matK (Hollingsworth et al., 2009).

Several groups of macroinvertebrates are frequently used to report EQS in the WFD. Species-level information on crustaceans, molluscs and the insect orders Ephemeroptera, Plecoptera and Trichoptera (EPT) are widely used. However higher taxa, e.g. genus- or family-level, are also used as BQEs and while some countries only use family-level identifications others use a mixed taxon approach, e.g. the River Invertebrate Classification Tool (<u>RICT</u>) (Davy-Bowker et al., 2008), used in the UK. There is a great variation between countries in which taxa are used to report to the WFD. For instance, freshwater assessments in the Netherlands utilize 224 species of the dipteran family Chironomidae when reporting water quality status, while Norway does not include species level information on any Diptera. This national-level taxonomic variation in part reflects the natural difference in species occurrences, but is necessary to consider when analysing gaps in the barcode libraries.

Freshwater fish are among the most commonly used organisms for assessing EQS according to the WFD, and their community composition and structure is the base for a high number of different metrics in Europe (Birk et al., 2012a). Sampling is conducted using a variety of methods, including electro-fishing or netting and should deliver data on abundance, species composition and age structure of fish present in a water body. However, large differences between countries exist in the percentage of occurring species considered for an assessment, and whether non-native species influence the overall score or not. In Ireland for example, all freshwater fishes are considered for WFD monitoring, while in Austria or Germany only about 60% of the complete fauna is routinely used. While according to practitioners, additional species encountered during sampling are often listed as an amendment to the official sampling protocols and reports, but they often have no impact on the BQE score because the species are not considered in the reference condition. Individual barcoding of sampled freshwater fish is of little use in biomonitoring of natural habitats. However, assessing and monitoring of freshwater fish diversity using eDNA from water followed by metabarcoding can be both more effective and more accurate than traditional specimen sampling (Hänfling et al., 2016; Valentini et al., 2015). Studies have indicated that the standard DNA barcode marker (COI) might not be optimal for this use (Kat Bruce & Emre Keskin pers. obs.), likely since non-target organisms are co-amplified with the available primers and mask the DNA signal from fish. Thus, a much higher sequencing depth is needed to reliably detect all fish species occurring in the studied waterbody, and constitutes suboptimal usage of available resources. Studies have shown that a hypervariable region of the rRNA 12S marker is a suitable target to amplify fish eDNA (Civade et al., 2016; Miya et al., 2015). As also discussed and successfully tested in DNAqua-Net WG3 (Field & Lab Protocols) this marker has a high potential to become the gold standard for regular eDNA-based fish monitoring in the future. We therefore also evaluate the completeness of the reference library for European freshwater fish species for 12S sequence data.

#### Aim of this study

The purpose of this paper is to identify gaps in DNA barcode reference libraries that are relevant for European countries when reporting water quality status to the EU in the context of the WFD and MSFD. The gaps for freshwater taxa are reported by country and taxonomic group, and compared across Europe, while gaps for marine organisms are evaluated by taxonomic group. We also discuss the necessity of both quality assurance and quality control (QA/QC) when building and curating a barcode reference library, and provide recommendations for filling the gaps in the barcode library of European aquatic taxa.

## 2. Material and methods

### 2.1 Checklists and datasets

Checklists of taxa used for freshwater EQS assessments according to the WFD were obtained from 30 nations (Supplement 1) through national contact points that were in direct contact with their countries’ environment agencies, water authorities, or water research institutes (see acknowledgements). National lists were sorted by taxon and assigned taxonomic coordinators among the authors who concatenated lists and unified the taxonomy (e.g. removing synonyms, checking validity of names, etc.) while keeping the country information for each taxon.

For marine species we used two generally accepted checklists to perform the gap-analysis of species relevant to the MSFD and WFD: AMBI - an index designed to establish ecological quality of European coasts, and ERMS (Costello, 2000). With the European focus of this analysis we delimited the AMBI list to a geographical selection by compiling only the species with European occurrence that include the following regions: Barents Sea, Norwegian Shelf, British Isles, Baltic Sea, North Sea, Celtic-Biscay Shelf, Iberian Coast, Mediterranean Sea, and Black Sea. The geographic distribution of each species on the original AMBI list was assessed through the World Register of Marine Species (<u>WoRMS</u>), as well as by the Ocean Biogeographic Information System (<u>OBIS</u>). The ERMS checklist on BOLD created by Dirk Steinke, titled ‘Marine Animals Europe’ (BOLD checklist code: CL-MARAE; last updated on 20^th^ March 2017), was used in this analysis. It contains records of 27,634 marine animals. A selection consisting of 21,828 species was used for further analysis, including taxonomic entities: Annelida, Arthropoda: Decapoda and Peracarida, Brachiopoda, Chordata: Euchordata - Pisces, Cnidaria, Echinodermata, Mollusca: Bivalvia and Gastropoda, Nemertea, Priapulida, and Sipuncula. We focused on benthic macroinvertebrates and fish and did not look specifically into meiofauna or pelagic animals (except fish), although many of the included species may have life-stages occurring in both environments.

Vascular plant checklists were checked for synonyms using three public databases: The International Plant Names Index (http://www.ipni.org), The Plant List (http://www.theplantlist.org) and Tropicos^®^ (http://www.tropicos.org).

For freshwater fish, we treated Europe as geographic entity, not by its political borders, but follow its definition as a “continent” with Turkey, Russia and Kazakhstan being only partly included and only with faunistic elements occurring in watersheds that lie within Europe (see also (Kottelat and Freyhof, 2007)). All lists were made available to taxonomic coordinators of selected taxonomic groups (specialists among the authors) to assure conformity of taxonomy and correct spelling. In this process, the taxonomic validation tool available from the Global Biodiversity Information Facility (<u>GBIF</u>), and WoRMS were used. For fish, the applied taxonomy mostly follows the international Catalog of Fishes (Fricke et al., 2018), which is also the backbone for the BOLD taxonomy.

Finalized species-level checklists were concatenated and uploaded to BOLD, and initial gap-analysis reports were retrieved. The reports were examined by taxonomic specialists to see if any reported gaps were due to taxonomic incongruence between the checklist and the BOLD taxonomic backbone. These were corrected in the uploaded checklists before final analysis (Supplement 2). Separate spreadsheets retaining the country information for each taxonomic group were kept for downstream analyses.

### 2.2 Gap-report analyses

Two sources of data were retained from BOLD for the majority of the taxonomic groups. Firstly, the checklist progress report option implemented in BOLD was used. Secondly, the checklists were compared to all publicly available sequence information in BOLD by using datasets for each taxonomic group. Progress reports and datasets were generated on the 6^th^ July 2018 for all groups except freshwater fish (1^st^ February 2018), freshwater Annelida (17^th^ September 2018) and Odonata (29^th^ November 2018). The dataset for Diptera used for the reverse taxonomy analysis was generated on the 18^th^ December 2018. The analyses were based on one or two barcode markers, depending on the taxonomic group (see Table 2).

Based on the BOLD gap reports, gap-analyses and summarizing statistics were calculated for all taxonomic groups using an analytical pipeline of custom made python scripts [deposited in GitHub https://github.com/dnaquanet/gap-analysis.git]. This pipeline was largely the same for all groups, except where specified under specific taxon treatment sections.

The data from taxonomic checklists with country information (i.e. nations in which the respective species are monitored) were combined with the information from BOLD. Species-based summaries were generated containing the number of countries in which a species is monitored by extracting the information from the taxonomic checklists. In addition, the total number of reference sequences stored in BOLD (i.e. sequences ≥ 500 bp), hereafter referred to as DNA barcodes, were taken from the progress report of each checklist. Additional BOLD quality criteria for barcodes, such as the availability of a trace sequence, were not considered. Using information from the publicly available data from the dataset output, it was possible to calculate the number of barcodes publicly stored in BOLD (BOLD public) or mined from GenBank (GenBank) as well as the number of privately stored barcodes in BOLD (BOLD private). Sequences flagged due to potential contamination, misidentification, or presence of stop-codons, were excluded from the analyses. For some species, DNA barcodes were deposited under the valid species name as well as under synonyms. In these cases, synonyms were part of the BOLD checklists and the barcode hits were merged to the valid species names.

In a further step, the proportion of species represented by a minimum number of DNA barcodes (threshold of 1 or 5) was calculated for each checklist. Additionally, country-based summaries were generated, providing an overview of the number of monitored species together with the percentage of barcode coverage for each taxonomic group in the reference libraries (threshold of 1 or 5). For both summary overviews, the available barcode information was sorted into three classes: BOLD public, BOLD total (including BOLD public and BOLD private) and total (including BOLD public, BOLD private, and GenBank). The data were visualized using the python-module matplotlib (Hunter, 2007) and cartopy (scitools.org.uk/cartopy) together with geographical information from naturalearthdata.com.

In contrast to all other gap-analyses, no geographical data were included for the marine taxa. Hence, the country-based analysis steps of the pipeline were omitted. Due to the large size of the ERMS checklists, no datasets could be produced in BOLD. Thus, only the results of the progress report were analysed for the availability of reference sequences. In the analysis of species used to calculate the AMBI, datasets could be produced in BOLD, and our analyses could distinguish between BOLD public, BOLD private, and GenBank sequence data.

To identify if species belonging to different ecological groups of the AMBI are equally well represented by reference sequences, a further gap-analysis was performed with species classified based on their ecological value.

For diatoms, the Diat.barcode library version 7 (Rimet et al., 2016) rather than BOLD was used, as this database is curated by diatom experts to ensure high-quality barcodes. Two genetic markers (rbcL and 18S) are used for barcoding diatoms (e.g. (Vasselon et al., 2018; Zimmermann et al., 2014), and the taxonomic checklists were compared to all available rbcL and 18S data in the database. Both, valid species names and synonyms were considered; subspecies were also accepted as valid. An overall gap-analysis and country-based summaries were generated. However, only a threshold of 1 was used. As all barcodes in Diat.barcode are publicly available at https://www6.inra.fr/carrtel-collection_eng/Barcoding-database, the differentiation between public and private data did not apply. Due to the high species diversity in diatoms, estimated at 100,000 (Mann and Vanormelingen, 2013), many low-frequency species could potentially negatively impact the barcode coverage, while the high-frequency (abundant) species could be sufficient for monitoring (Lavoie et al., 2009). Hence, we re-analysed the barcode coverage for two checklists (France freshwater phytobenthos and Croatia marine diatoms) using only high-frequency species.

Two standard barcode markers (rbcL or matK) are accepted for vascular plants in BOLD. However, the checklist progress report does not include information on which of the two barcode markers were covered for each taxon. Hence, the first part of the analyses described above was conducted for vascular plants regardless of which of the two markers was present (rbcL OR matK). In contrast, the BOLD dataset includes information on which marker is sequenced for a certain record. Hence, for the public data (BOLD public and GenBank) gap-analyses were performed for each marker as well as for the combination of both markers (rbcL AND matK).

For gap-analysis of freshwater fish we also included the 12S marker. Since there are no 12S sequence data available in BOLD (as of February 1^st^ 2018) for European freshwater fishes, we manually compared our target species list with the available mitochondrial genomes from MitoFish (http://mitofish.aori.u-tokyo.ac.jp), and NCBI’s RefSeq and Nucleotide databases. All available sequence data for Actinopterygii (whole mitochondrial genomes and full or partial 12S sequences) were imported into the software Geneious version 7.1.9 (Biomatters Ltd, New Zealand) and after aligning with the MAFFT-plugin (Katoh and Standley, 2013) trimmed to the hypervariable region of the 12S rRNA gene using the published primer pair MiFish-U/E (Miya et al., 2015) as correctly given in Ushio et al. (Ushio et al., 2018). In the final alignment only species present with sequence information for this locus (ca. 175 bp) were retained and used for the gap list evaluation. Due to the completeness of the barcoding databases for species used in country-based monitoring lists, in general, no geographical information was used for the gap-analysis. However, a map was generated for species of the European-wide fish list where barcodes are still missing.

Finally, we refrained from providing any particular DNA barcode gap-analysis for groundwater ecosystems and their species pools. This is because the biological component is currently not considered for subterranean freshwater monitoring and reporting under the umbrella of the WFD, which relies on the chemical status and water quantity in aquifers instead.

### 2.3 Reverse taxonomy

As a case study, we analysed the proportion of public barcodes originating from reverse taxonomy for freshwater macroinvertebrates, i.e. specimen identification via its DNA barcode and not by morphology. In the datasets obtained from BOLD, the entry “Identification Method” was screened for the presence of several keywords e.g. “BOLD ID Engine”, “BIN Taxonomy Match”, “Tree based identification” or “DNA Barcoding”. A full list is deposited in Supplement 3. For each species, the number of public barcodes originating from reverse taxonomy was compared to the total number of available public barcodes in BOLD. Four cases were considered, in which reverse taxonomy can have a strong influence: i) all public data originates from reverse taxonomy, ii) more than half of the public data originates from reverse taxonomy, iii) only when including barcodes based on reverse taxonomy, at least five public barcodes are present and iv) when less than five public barcodes are present, at least one originates from reverse taxonomy.

## 3. Results

Our results revealed considerable variation in barcode coverage for selected major groups in the queried databases (Table 1). Freshwater vascular plants and freshwater fish had the largest coverage, though still less than 70% of the species had five or more barcodes available. The lowest barcode coverage is found in the marine invertebrates of the ERMS list 9.9% (five or more barcodes) to 22.1% (one or more barcodes) and diatoms (14.6%), while more than 60% of the 4502 freshwater invertebrate species used in ecological quality assessments of freshwater ecosystems had one or more barcodes (Table 1).

**Table 1.**
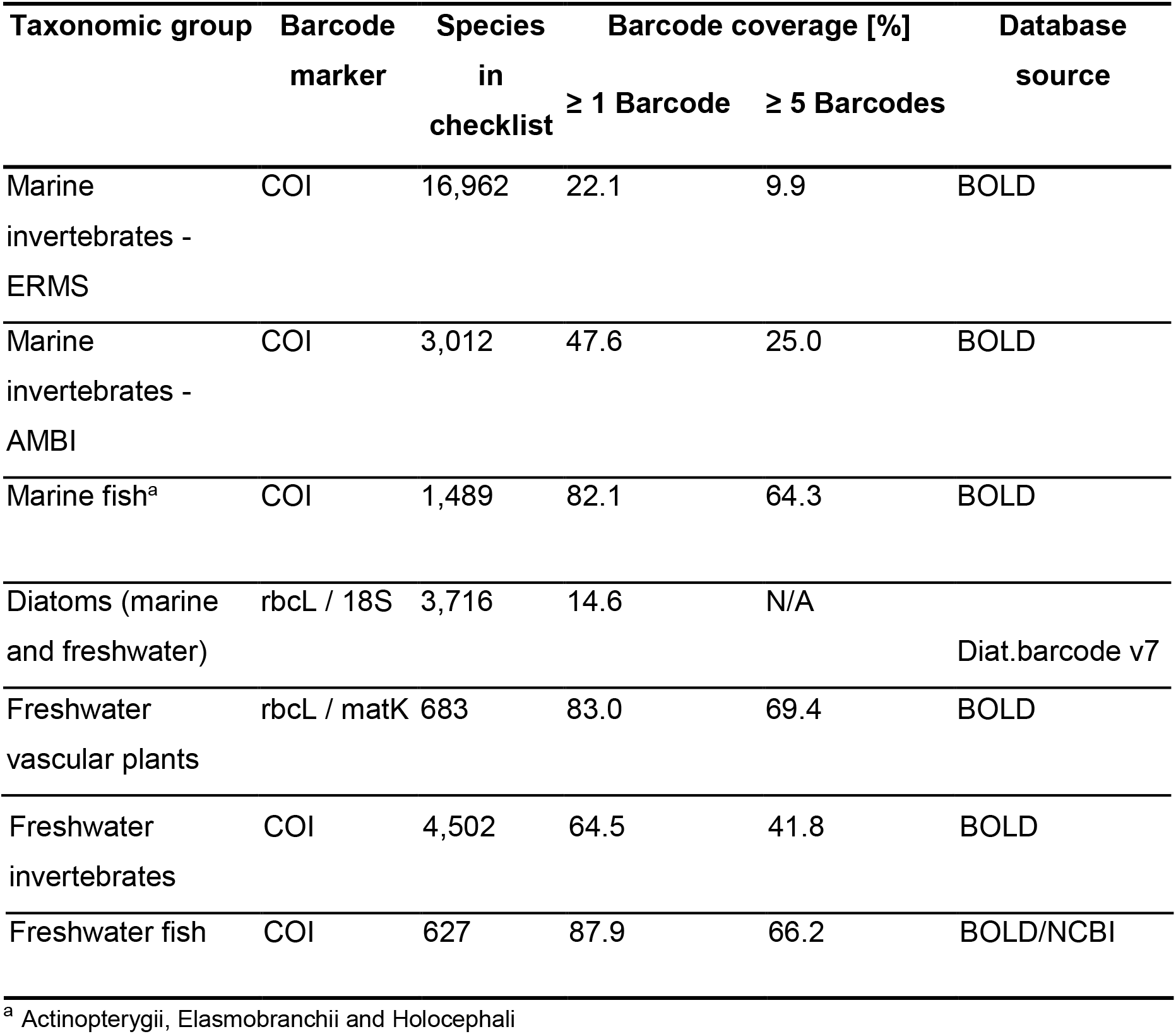
Overall barcode coverage for selected major groups.

### 3.3 Marine macroinvertebrates & fish

#### 3.3.1 Gap-analysis for the European AMBI-list

A total of 3,012 marine species were compiled in the AMBI checklist for Europe. Forty-eight percent of them have at least one representative DNA barcode sequence in either BOLD or GenBank, but as much as 23% of those species only have private records (Fig. 1, Supplement 2), and 22% of those with barcodes are single specimen records.

**Figure 1.**
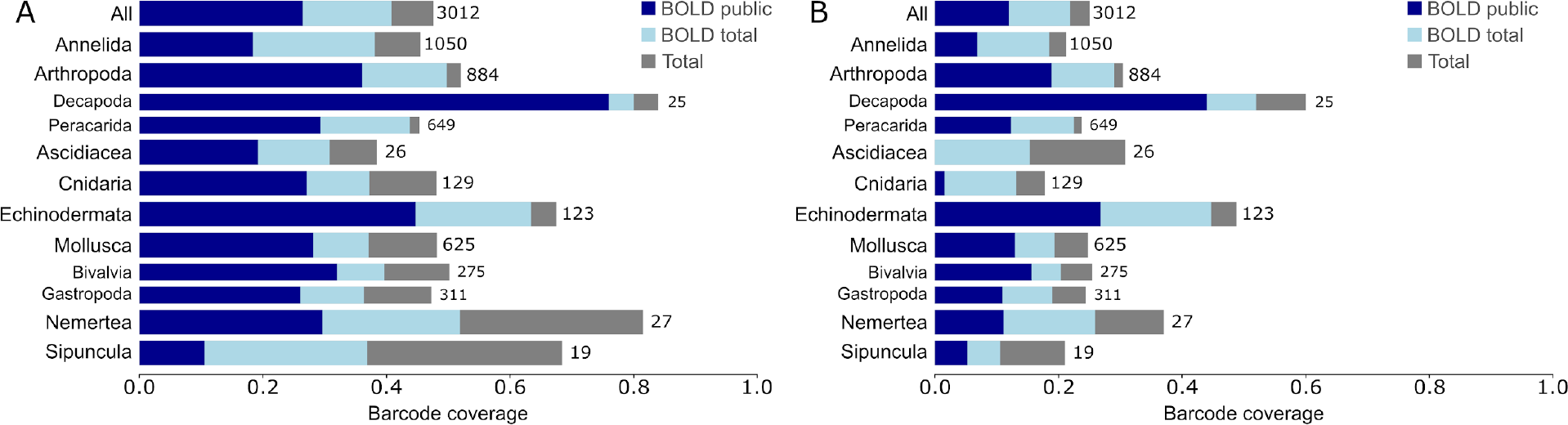
Cumulative barcode coverage for marine invertebrates in the AMBI list. Barcode coverage of at least 1 reference sequence (A) or a minimum of five reference sequences (B). If barcodes of a species were not recorded in the BOLD public library, the BOLD private library was queried, and subsequently GenBank. Numbers on bars refer to total number of species in checklist. Thick bars represent phyla, thin bars represent taxa of lower taxonomic rank. Taxonomic groups with less than ten species are not indicated.

Among the 10 largest taxonomic groups included in this particular analysis, the Chordata (excluding Vertebrata) displayed the lowest proportion of species with DNA barcodes (38%), though only 26 species (within Ascidiacea) were listed for this taxon. In comparison, the best represented taxon was the Nemertea, which has DNA barcodes for 81% of the 27 species considered, while the second most complete group has 67% (Echinodermata). Most of the remaining taxa have completion levels between 40 and 50%, including the three most species-rich taxa (Annelida, Mollusca and Arthropoda), that comprise 85% of the species in the European AMBI checklist (Fig. 1).

A narrower analysis of Mollusca shows that Bivalvia and Gastropoda have only moderate levels of completion (50 and 47%, respectively), whereas within malacostracan crustaceans, Decapoda (Arthropoda) is far more complete (84%) than Peracarida (45%). However, the number of species considered is highly disparate for these two groups (25 Decapoda vs. 649 Peracarida) (Fig. 1). The proportion of singletons (i.e. only one barcode sequence available) per taxonomic group ranges from 10% to 25%, although for some taxa the observed proportion of singletons was considerably higher (e.g. 50% in Brachiopoda and 38% in Sipuncula).

Most of the species from the AMBI checklist have public DNA barcodes available either from BOLD or GenBank, with only 11% represented exclusively by private records. Two groups have slightly higher values, Echinodermata (15%) and Arthropoda (12%). The levels of completion by AMBI’s Ecological groups (I to IV) are similar, ranging from 43% in group IV to 56% in group III (Supp. Fig. 1). However, 215 species were not assigned to ecological groups, and among these the completion is low (ca. 38%). Species barcodes found exclusively in BOLD private range from 10% (IV) to 13% (V) in each of AMBI’s ecological groups.

#### 3.1.2 Gap-analysis for the ERMS checklist

The selection from the ERMS list on BOLD contains 16,962 species. Twenty-two percent of these species have at least one DNA barcode in BOLD (Fig. 2). Of these species, 26% have singletons and nearly 10% have five or more DNA barcodes. These figures include DNA barcodes from GenBank that are present in BOLD. The highest coverage is found in Decapoda (50%), followed by Sipuncula (42%), a phylum with 45 species only found in the ERMS list (Fig. 2). At the other end, the lowest coverage (11%) is observed in Brachiopoda (37 species). Nemertea also have a low coverage, 15% for the 380 listed species. The coverage of most other taxonomic groups ranges from 20 to 30%.

**Figure 2.**
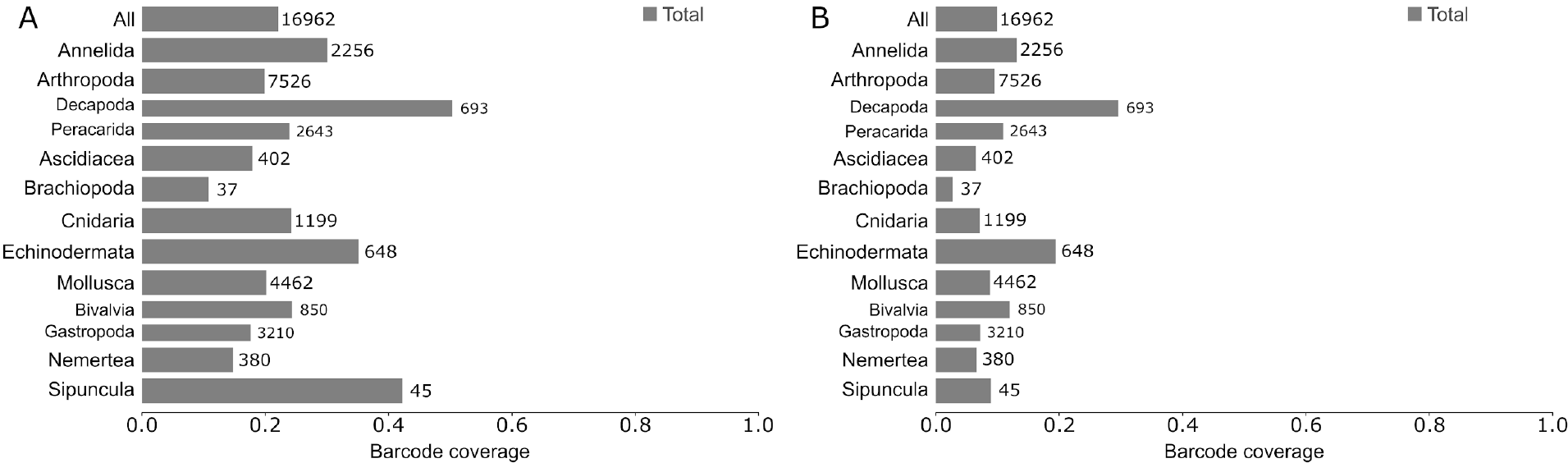
Barcode coverage for marine invertebrates of the ERMS checklist. Barcode coverage of at least 1 reference sequence (A) or five reference sequences (B). Thick bars represent phyla, thin bars represent taxa of lower taxonomic rank. Numbers on bars refer to total number of species in checklist. Taxonomic groups with less than ten species are not indicated.

Within phyla, there are clear differences in the proportion of DNA barcodes between taxonomic subgroups. Arthropods have a coverage of 20% as a whole, but the Decapoda reach 50%, while the Peracarida reach only 23%. Within Mollusca, with an overall coverage of 20%, Bivalvia reach 24% and Gastropoda 18%. The proportion of singletons roughly follows the inverse pattern as the proportion of total DNA barcodes: the lowest proportion of 8% is found in marine fish, while the highest proportion of 57% is found in Brachiopoda.

A detailed analysis of cnidarians in the ERMS checklist reveals that while 353 of the 1,201 species (29.4%) are listed with sequence information in BOLD, only 97 species (8.1%) have sequences that meet the formal barcode requirements. We observed that many of the sequences were mined from GenBank, containing limited information and are a potential source of errors. A similar situation was observed for ascidians where 84 out of 402 species in the ERMS checklist (20.9%) have sequence information while, only 5.7% of the species had references to vouchers and sufficient metadata to be barcode compliant.

The marine fish checklist obtained from ERMS includes 1,489 species partitioned among the three most prominent classes examined as follows: Actinopterygii (1,339), Elasmobranchii (143) and Holocephali (7). Overall, 82% of the species are barcoded (64% ≥ 5 barcodes), ranging from 100% (71% ≥ 5 barcodes) for the Holocephali to 81% (63% ≥ 5 barcodes) for the Actinopterygii, with the Elasmobranchii coverage is in between (92% ≥ 1 barcodes, 80% ≥ 5 barcodes) (Fig. 3).

**Figure 3.**
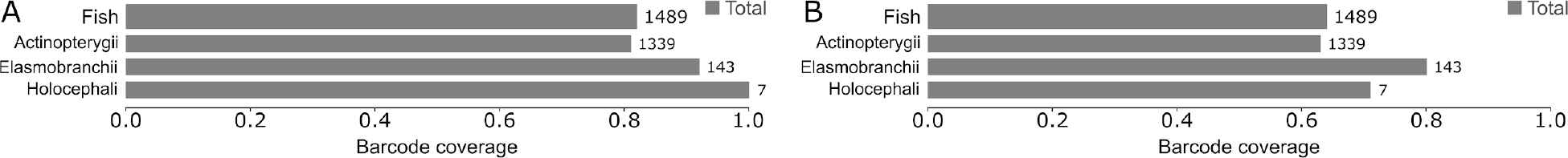
Barcode coverage for marine fish of the ERMS checklist. Barcode coverage by at least 1 reference sequence (A) or five reference sequences (B). Thick bars represent all fish, thin bars represent lower taxonomic rank. Numbers on bars refer to total number of species in checklist.

### 3.2 Diatoms

Taxonomic checklists for diatoms were obtained from 16 countries and contained a total of 3,716 species ranging from 6 (Albania) to 2,236 species (France). This list covers very different habitats, freshwater phytobenthos, freshwater phytoplankton and marine phytoplankton. Some national checklists did not mention which habitat was covered.

The general coverage of diatoms was very low, with 15% of all species having at least one sequence of rbcL or 18S (Fig. 4). The coverage of rbcL (13%) is slightly better than the coverage of 18S (11%). However, in most cases both markers are present if any sequence is available (9%). Per country, the coverage ranged from 10% (France) to 37% (Italy), when both markers are present and 15% (France) to 55% (Italy), when at least one of the markers is present (Suppl. Fig. 1).

**Figure 4.**
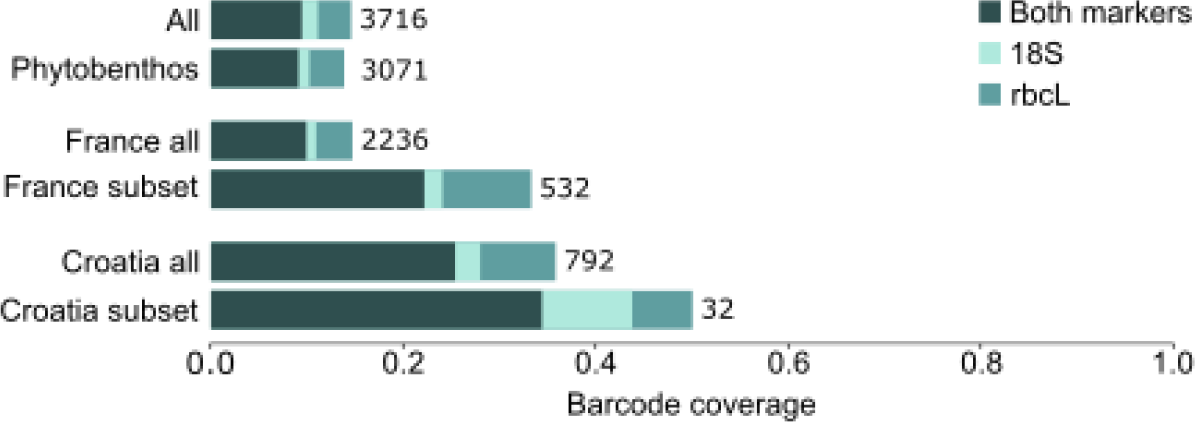
Cumulative barcode coverage of diatoms.

A gap-analysis of diatoms ranked by the number of countries that monitor those species, revealed that the most frequently monitored species have a moderate to high representation for both markers (Fig. 5A). For the 16 species used in 14 countries, 81% have rbcL and 18S data and additionally 13% have rbcL data only. For species monitored by few countries, the barcode coverage is comparatively poor (below 20% for species monitored in ≤ 7 countries).

**Figure 5.**
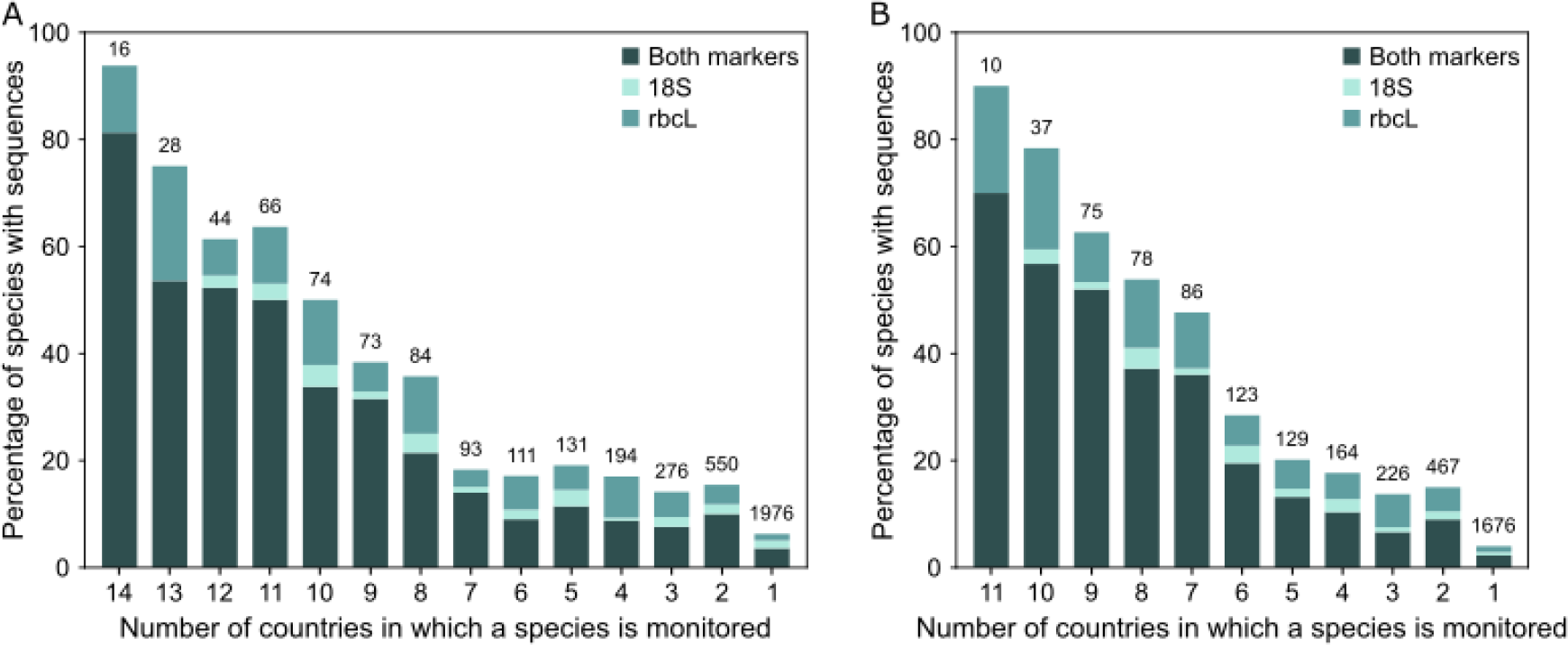
Cumulative barcode coverage of diatoms by the number of countries monitoring them. (A) All diatom species, (B) freshwater phytobenthos species. Numbers on bars refer to the number of species per country category.

For freshwater phytobenthos, the habitat where diatoms are used most often as ecological indicators, the most frequently monitored species have a moderate to high representation of both markers (Fig. 5B). Similar to all diatom datasets, most of the species monitored in eleven countries are represented by both markers (70%), with additional species barcodes for rbcL (20%). For species monitored by fewer countries, the coverage is considerably smaller (below 20%, for species in ≤ 4 countries).

For the most common species of freshwater phytobenthos monitored in France, 553 of the 2,236 species were scored as abundant. In this subset, the barcode coverage was 33%, considerably higher than the 15% of all species. The proportion of species with both rbcL and 18S sequenced was 20% compared to 10% for all species (Fig. 4). A similar picture was evident for the marine diatoms from Croatia. Of the 100 most frequently observed marine phytoplankton species (including Diatoms, Dinoflagellates, Silicoflagellates and Coccolithophorids), 32 were diatoms. Of these 32 species, 50% had at least one barcode available compared to 36% in the total dataset of 729 species. The proportion of species with both barcodes was 34%, compared to 25% for all species (Fig. 4).

### 3.3 Vascular plants

General taxonomic checklists for freshwater macrophytes were obtained for 16 nations. The compiled list of 1,242 species names was filtered for vascular plants, resulting in 683 species. In general, vascular plants are well covered by one or the other standard barcode marker, with more than 83% of the species having at least one sequence, and 69% having at least 5 sequences (Tab. 1).

Compared with public records, however, these results seem slightly overemphasized as 22% (153) of the species have no rbcL nor matK sequences publicly available on BOLD (or mined from GenBank, Fig. 6A). Moreover, only 46% (316) of the species have barcodes for both loci publicly deposited in BOLD. The remaining 214 species have incomplete data: *i*) rbcL publicly deposited in BOLD, but matK sequences absent (53), or mined from GenBank (80); ii) sequences for both loci coming from GenBank (38); *iii*) sequences for only one locus issued from GenBank (rbcL - 28; matK - 15).

**Figure 6.**
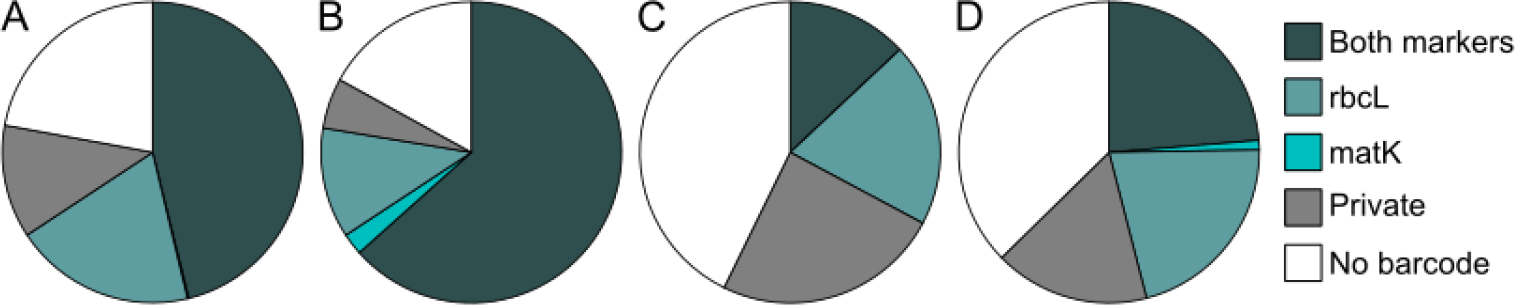
Barcode coverage for freshwater vascular plants. (A) ≥1 DNA barcode available in BOLD, (B) ≥1 DNA barcode available in BOLD or GenBank, (C) ≥5 DNA barcodes available in BOLD or (D) ≥5 DNA barcodes available in BOLD or GenBank.

In sum, *rbcL* is the best represented DNA barcode marker for vascular plants with 75% of the species having publicly deposited sequences, and 66% of the species having BOLD public data (Fig. 6). Sixty-six percent of the species have publicly deposited barcodes for matK, with only 46% of the species having sequences deposited in BOLD public.

The number of monitored species varied strongly, ranging from six (Poland) to 394 (Hungary, Fig. 7A). The average barcode coverage (BOLD and GenBank data) was relatively evenly distributed with a minimum of 76% (Lithuania), reaching 100% in three countries (Austria, Poland and Switzerland, Fig. 7B). A higher and more homogeneous coverage was found for rbcL (67 - 90%; Fig.7C) than matK (0 - 74%; Fig. 7D), both for BOLD public and GenBank data (rbcL: 71% - 100%; matK: 50% - 87%; Supp. Fig. 2). Two species were monitored in twelve countries (*Alisma lanceolatum* and *A. plantago-aquatica*) and approximately one fifth of the species in more than 4 countries (Fig. 7E, F). The barcode coverage of these species was 100% when public and private data were taken into account. It decreased slightly for species monitored in four or fewer countries. Nevertheless, more than 40% of the 330 species monitored in one country only had rbcL and matK data deposited publicly in BOLD and 73 % had associated sequences when private BOLD and GenBank data were included.

**Figure 7.**
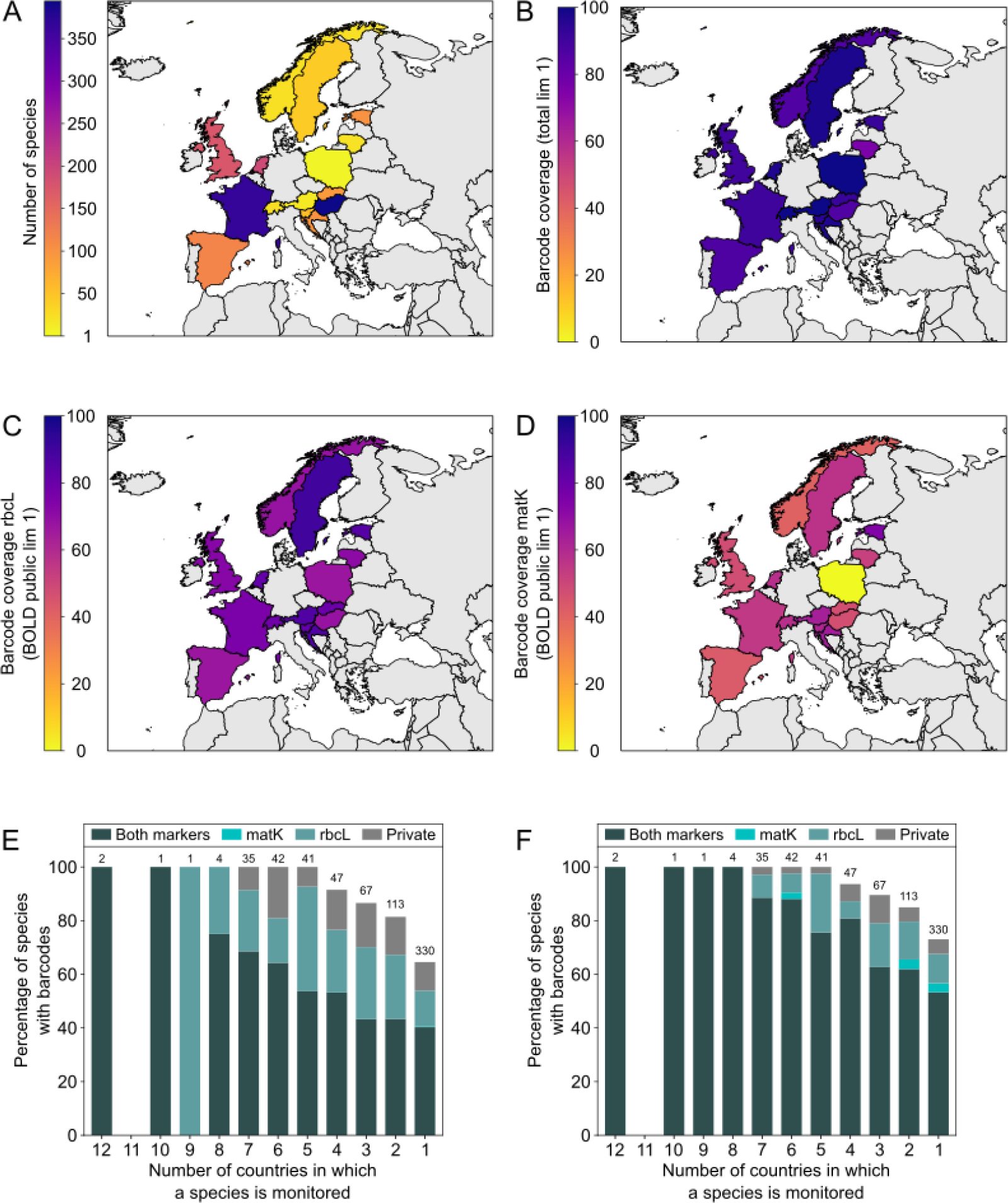
Barcode coverage maps for freshwater vascular plants (lim 1 = minimum one record). (A) Number of monitored species per country, (B) barcode coverage per country with all available data (total), (C) rbcL-specific coverage per country publicly available in BOLD, (D= matK-specific coverage per country publicly available in BOLD, (E) cumulative barcode coverage of vascular plants available in BOLD by number of countries monitoring them, (F) cumulative barcode coverage of vascular plants available in BOLD or GenBank by number of countries monitoring them.

### 3.4 Freshwater macroinvertebrates

The analysed national monitoring checklists comprise 4,504 species of freshwater macroinvertebrates, including insects (ca. 80% of the listed species), annelids (ca. 6%), arachnids (ca. 5%), crustaceans (ca. 4%), molluscs (ca. 4%), flatworms (ca. 1%) and nematodes (< 0.05%). When considering all species with at least one barcode in BOLD, 64.5% of the species are covered (Fig. 8). Most barcodes are publicly available. For the more strict criterion of ≥ 5 barcodes per species, only 41.9% of the species are covered. Among all taxonomic groups considered in the analysis, the three insect orders Odonata, Trichoptera and Hemiptera along with crustaceans are best covered with ≥80% of species barcoded from each taxonomic group. The groups with the least coverage are flatworms (less than 5%), followed by annelids, molluscs and certain insect orders, such as Diptera and Ephemeroptera, in which less than 60% of listed species are represented by at least one barcode (Fig. 8). Only in the case of Hemiptera, more than 80% of the species are represented by at least five barcodes while, except for Odonata, Trichoptera, Coleoptera and Crustacea, less than 50% of the species are covered in the other macroinvertebrate groups. For some groups, such as molluscs, annelids and crustaceans, a substantial share of the available reference sequences are not deposited in BOLD, but present in GenBank (Fig. 8). The most monitoring-relevant insect taxon with lowest coverage on BOLD is Diptera (ca. 60% of the 2,108 species in the list). Hemiptera, with 76 species listed and ca. 92% already barcoded will probably be the first group to have full coverage in the near future.

**Figure 8.**
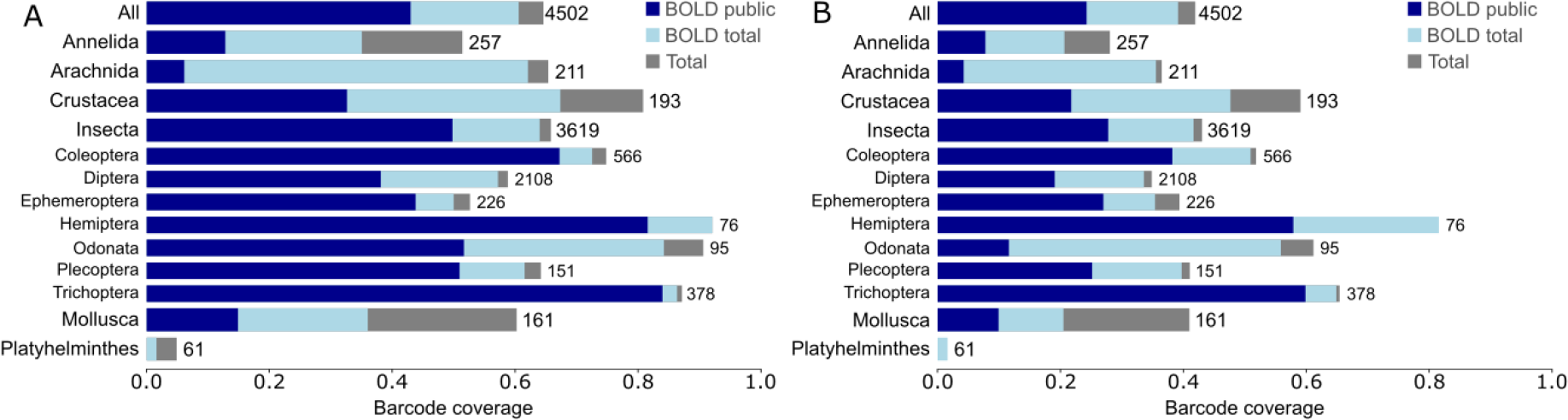
Cumulative barcode coverage for freshwater invertebrates. Barcode coverage by at least one reference sequence (A) or five reference sequences (B). If barcodes of a species were not recorded in the BOLD public library, the BOLD private library was queried, and subsequently GenBank.Thick bars represent higher taxonomic ranks, thin bars represent insect orders. Numbers on bars refer to total number of species in checklist. Taxonomic groups with less than ten species are not indicated.

#### 3.4.1 Insects

Insects are used for monitoring ecological status in 29 out of the 30 surveyed countries. All national monitoring checklists combined comprised 3,619 insect species (Supplement 2, Fig. 9D). However, taxonomic resolution used between countries differed substantially. Seven countries exclusively assess taxonomic groups above species level, two countries only above genus level, and five countries only above family level (Supplement 1). Assessed taxa per country range from 10 (Albania) to 2,903 (Czech Republic, Fig. 10). In total, eleven insect orders are monitored, ranging from orders with only one relevant species (Hymenoptera) to orders with 2,108 species (Diptera, Fig. 8). The top ten species monitored in most countries all belong to Ephemeroptera with *Ephemera danica* and *Serratella ignita* being the most frequently listed species (20 countries each).

**Figure 9.**
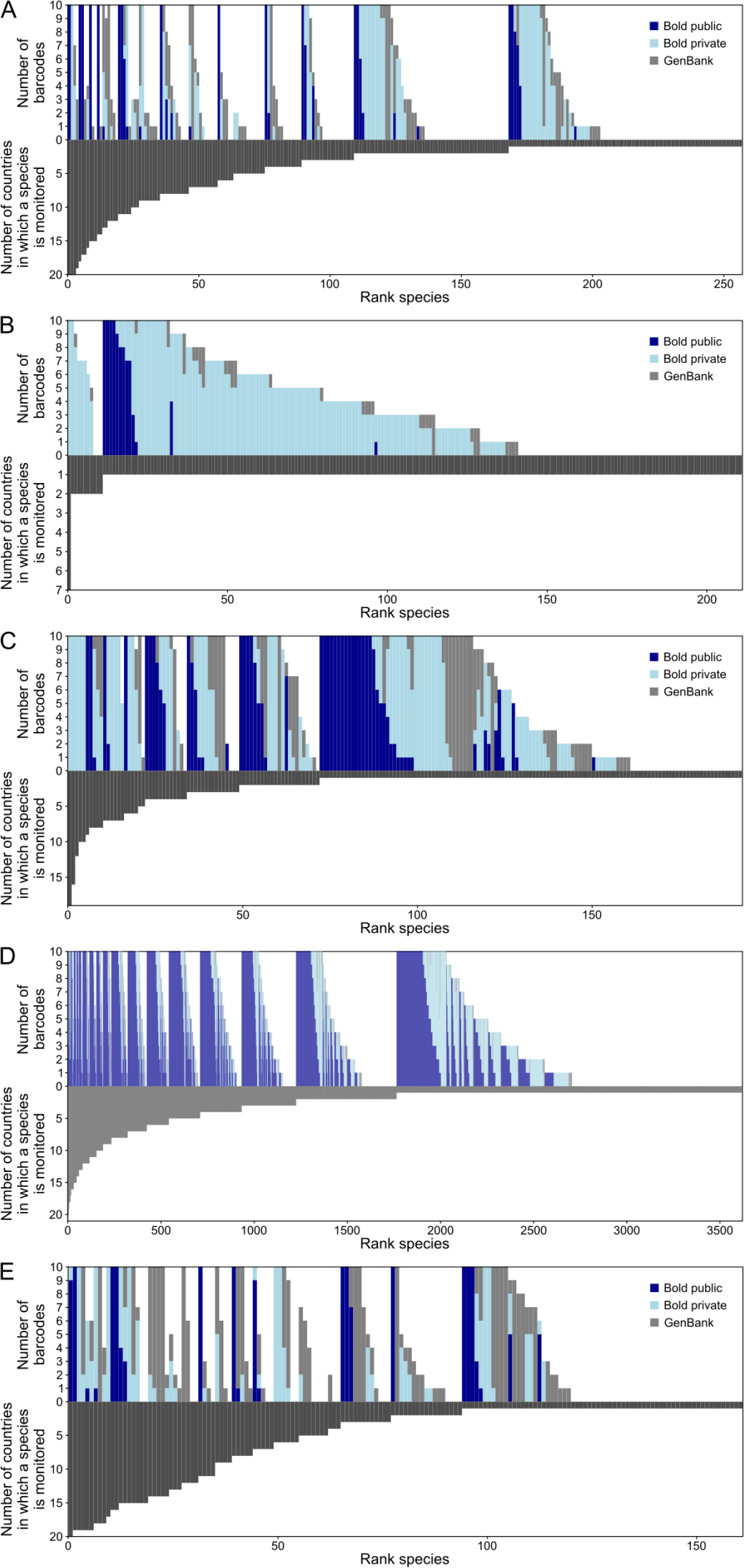
Barcode coverage per species. The upper bars show the barcode coverage (up to a maximum of 10 barcodes). The lower bars show the number of countries in which a species is monitored. (A) Annelida, (B) Arachnida, (C) Crustacea, (D) Insecta, (E) Mollusca.

**Figure 10.**
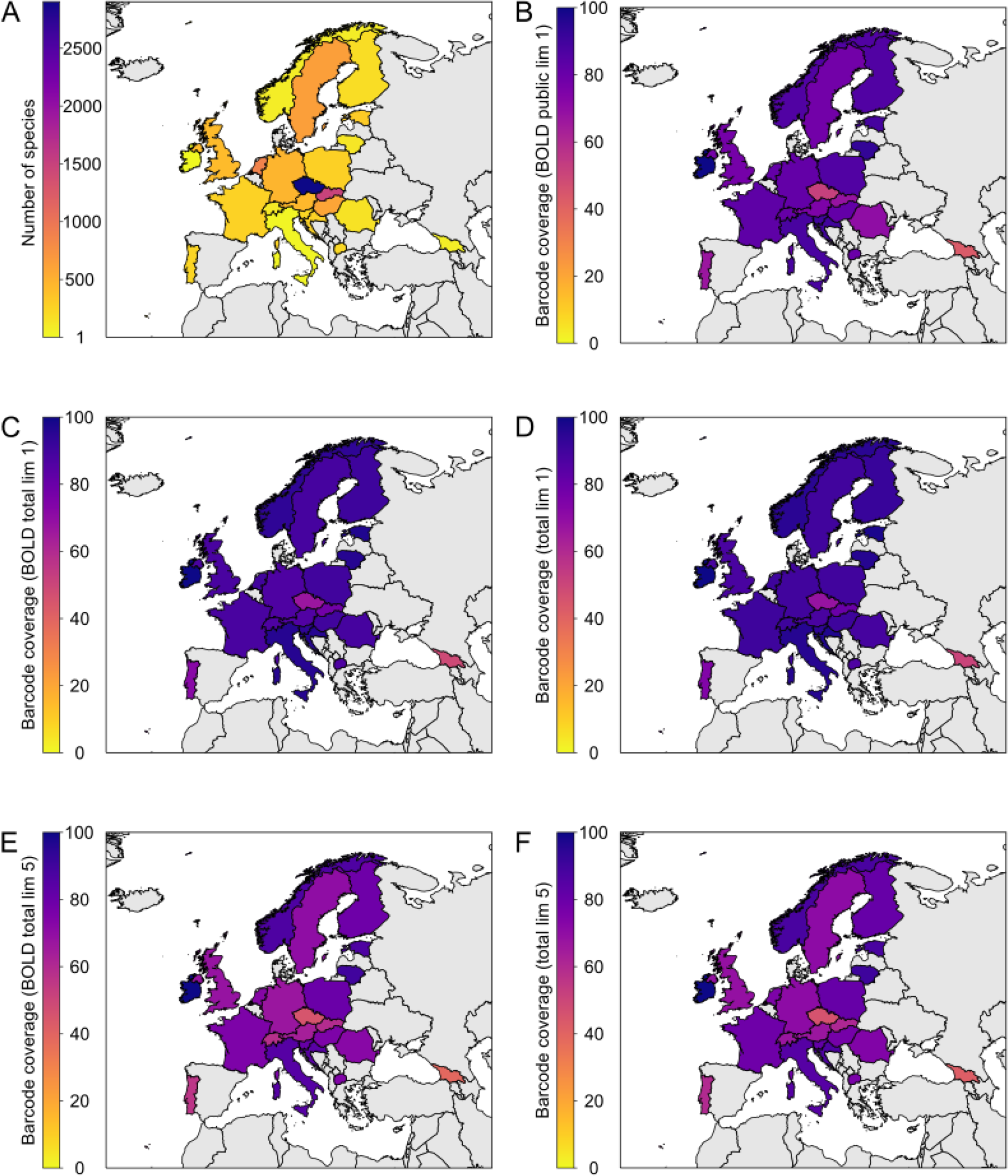
Barcode coverage maps of Insecta. (A) Number of monitored species per country. (B) - (F) Barcode coverage per country for different datasets (BOLD public, BOLD total and total) and thresholds (lim 1 = minimum one record; lim 5 = minimum five records).

On average, 65.7% of all monitored insect species are barcoded, ranging from orders with only 52.7% and 58.8% barcoded species (Ephemeroptera and Diptera) to highly covered orders (Trichoptera, 87.0%; Odonata, 90.5%; Hemiptera 92.1%; Fig. 8). A high proportion of barcodes for these species is deposited in BOLD (95.3%; 91,066 barcodes) of which 70.9% have publicly available metadata. However, for 513 barcoded species (14.2%) there is no BOLD public data. For the most frequently monitored species, *Ephemera danica,* there are only 4 public (and 11 private) COI barcodes in BOLD. In contrast to the top monitored species, 9 of the 10 species with the most barcodes (BOLD and Genbank combined) belong to Diptera with the two Chironomidae species *Paraphaenocladius impensus* (5,981 barcodes) and *Paratanytarsus laccophilus* (4,058) being the most often barcoded species. Of the 1,240 missing insect species that are monitored in at least one country, 917 are monitored in a single country (Czech Republic), and 674 of those species are exclusively monitored in that country. The coverage of barcoded species per country is on average 87.6%, ranging from 51% (Georgia, only mayflies) and 68% (Czech Republic) to 98% (Estonia) and 100% (Ireland, just three taxa monitored at the species level).

#### 3.4.2 Arachnids

A large proportion of the arachnid species records in BOLD is private (Fig. 8). The coverage of the 211 species reported from all countries in total is moderate with 65% of the species represented with at least one barcode. It is remarkable that 201 of the 211 arachnid species are only monitored by one country, the Netherlands (Fig. 9B). Of these, 200 are solely monitored in this country. The spider *Argyroneta aquatica,* which is monitored by the most countries (7), has only private reference barcodes in BOLD, and five sequences in GenBank.

#### 3.4.3 Crustaceans

A total of 193 crustacean species are included in the nationwide checklists; 22 of the 30 surveyed countries monitor one or more crustacean species (Fig. 11). They represent four classes: Branchiopoda (62 spp.), Hexanauplia (25 spp.), Ichthyostraca (3 spp.) and Malacostraca (110 spp.). Among them, the most frequently monitored species are the malacostracans: common waterlouse, *Asellus aquaticus*, monitored in 19 countries, the noble crayfish, *Astacus astacus* - in 16 countries, and the amphipod *Gammarus roeselii* - in 12 countries. Each of these species is covered with numerous private only reference barcodes in BOLD or publicly available sequences in GenBank (Fig. 9C). Thirty six species of crustaceans have no barcode coverage at all, neither in BOLD nor in GenBank, while 26 are covered only in GenBank. Among those covered in BOLD, 67 species are represented by private reference barcodes only. Most of the species (121 spp.) are monitored only in one country. For example, 53 species, predominantly branchiopods and hexanauplians, are monitored in Sweden only. Eleven of these species have no barcode coverage, neither in BOLD nor in GenBank, while 22 species are represented only by private barcodes. In general, the barcode coverage (including GenBank data) per country is good and relatively evenly distributed, from 70% to 100% of species barcoded in each country (Fig. 11D). These values drop down immensely when only the public BOLD data are taken into account (Fig. 11B). In the countries such as Italy and Ireland not even 10% species is covered, while only in Germany, UK, the Netherlands and Norway the coverage approaches 50% of the species monitored in each of these countries.

**Figure 11.**
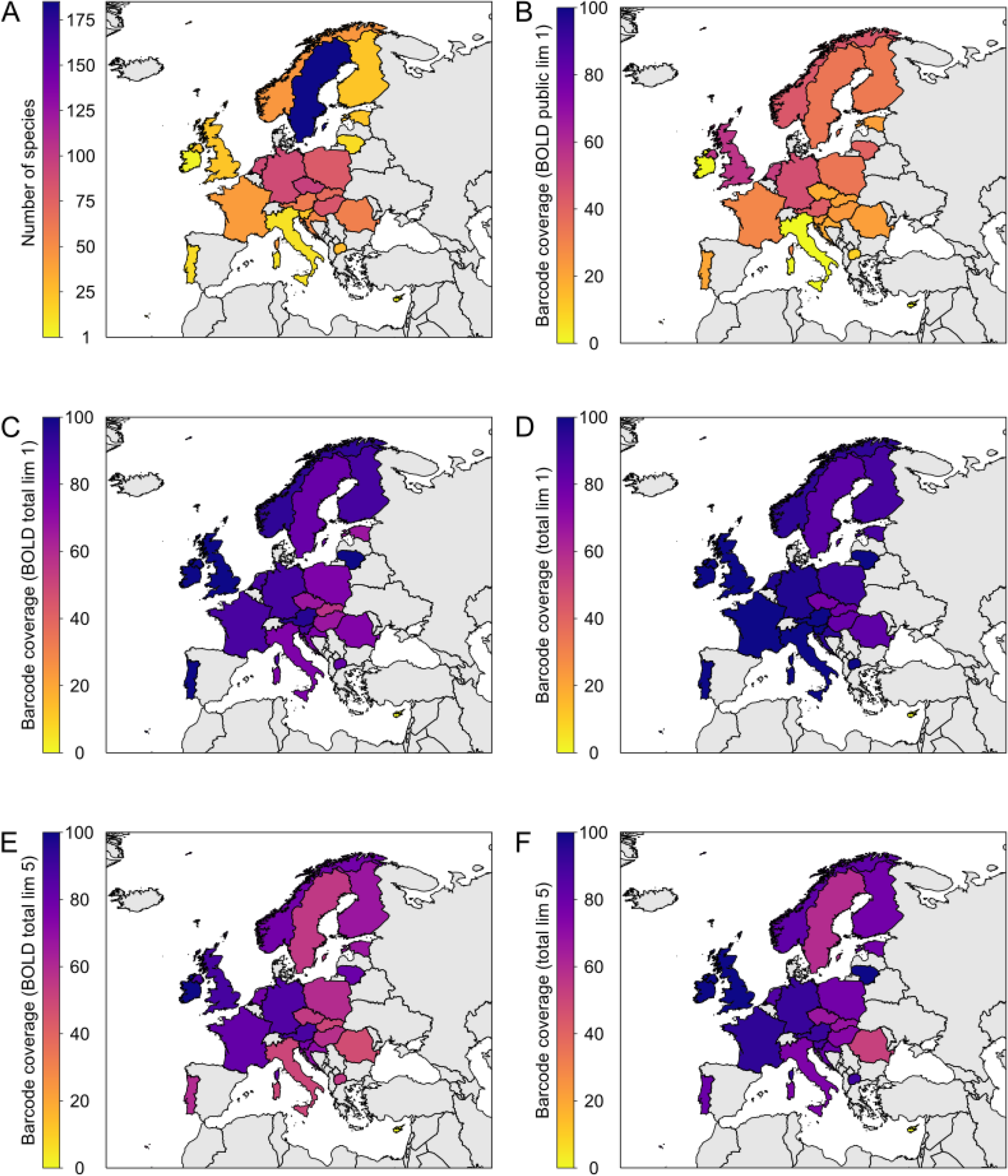
Barcode coverage maps of Crustacea. (A) Number of monitored species per country. (B) - (F) Barcode coverage per country for different datasets (BOLD public, BOLD total and total) and thresholds (lim 1 = minimum one record; lim 5 = minimum five records).

#### 3.4.4 Annelids

In total, 257 species of annelids are used in freshwater biomonitoring in the 21 countries that supplied lists (Fig. 12). They represent two classes, Clitellata with the subclasses of Oligochaeta, Hirudinea (leeches) and Branchiobdellida and Polychaeta with the subclass Sedentaria. Among them, three species of leeches, *Erpobdella octoculata*, *Glossiphonia complanata* and *Helobdella stagnalis* are monitored in 20 countries (Fig. 9A). Further 21 species of both leeches and oligochaetes are monitored in 11 to 19 countries. The most commonly monitored polychaete is the freshwater alien *Hypania invalida* included in lists of five countries. The other alien species, *Marenzelleria neglecta*, is generally brackish water and is monitored only in Germany. A couple of other brackish water native species are generally monitored in single countries only. Altogether almost 50% of the listed species are represented by DNA sequences. However, they are generally poorly represented in BOLD, where barcodes for ca. 40% of the species are deposited and only some 20% are publicly available. Most of the species with no barcodes at all are monitored in few countries only (predominantly in Czech Republic and Slovakia) with some notable exceptions, such as the leech *Alboglossiphonia heteroclita* present on the lists of 15 countries, and the oligochaete *Haplotaxis gordioides* monitored in 12 countries.

**Figure 12.**
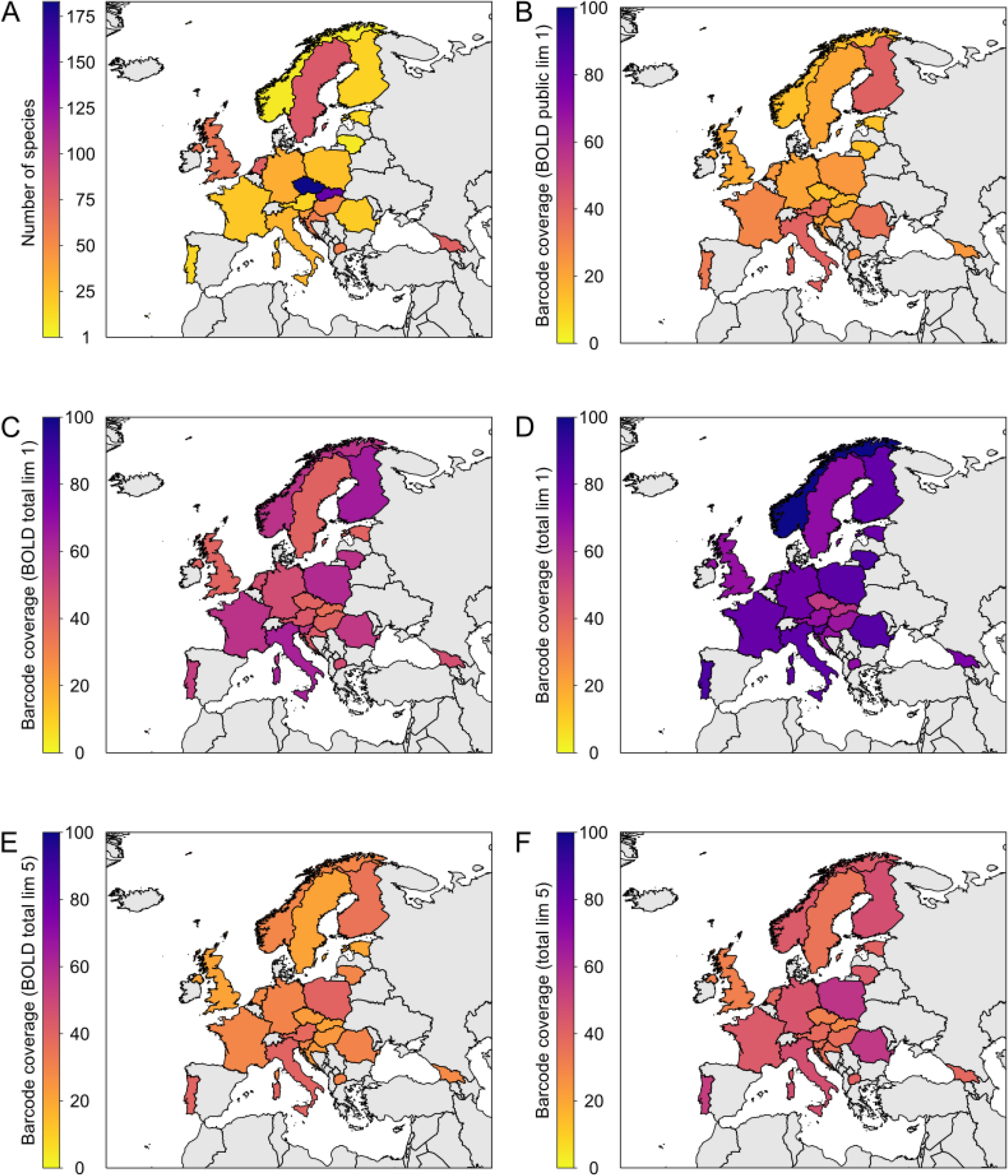
Barcode coverage maps Annelida (A) Number of monitored species per country. (B) - (F) Barcode coverage per country for different datasets (BOLD public, BOLD total and total) and thresholds (lim 1 = minimum one record; lim 5 = minimum five records).

Country wise, the barcode coverage (including GenBank data) extends from ca. 50% of species barcoded in Czech Republic and Slovakia to 100% in Norway (Fig. 12D). When only public BOLD records are considered, the barcode coverage per country drops down to 20%-40% (Fig. 12B).

#### 3.4.5 Molluscs

The national checklists of freshwater molluscs contain a total of 161 species, ranging from one (Cyprus) to 77 (Czech Republic) species per country (Fig. 13). *Ancylus fluviatilis*, the most commonly surveyed species, is included in 20 national checklists, while a total of 67 species are considered by a single checklist only (22 of them in Georgia) (Fig. 9D). The total barcode coverage of freshwater molluscs (about 60%) was in the range of most freshwater invertebrate groups (Fig. 8). While the proportion of species with public barcodes deposited in BOLD was relatively low (only 15%), the proportion of species with sequences derived only from GenBank was considerably high (24%). A similar pattern was evident when a minimum coverage of five barcodes was used (Fig. 8B). Here, 41% of the species met the criteria when all public and private data were considered, 10% of the species were covered in the BOLD public database, while 21% of the species only had sufficient barcodes if GenBank data were considered together with data from BOLD.

**Figure 13.**
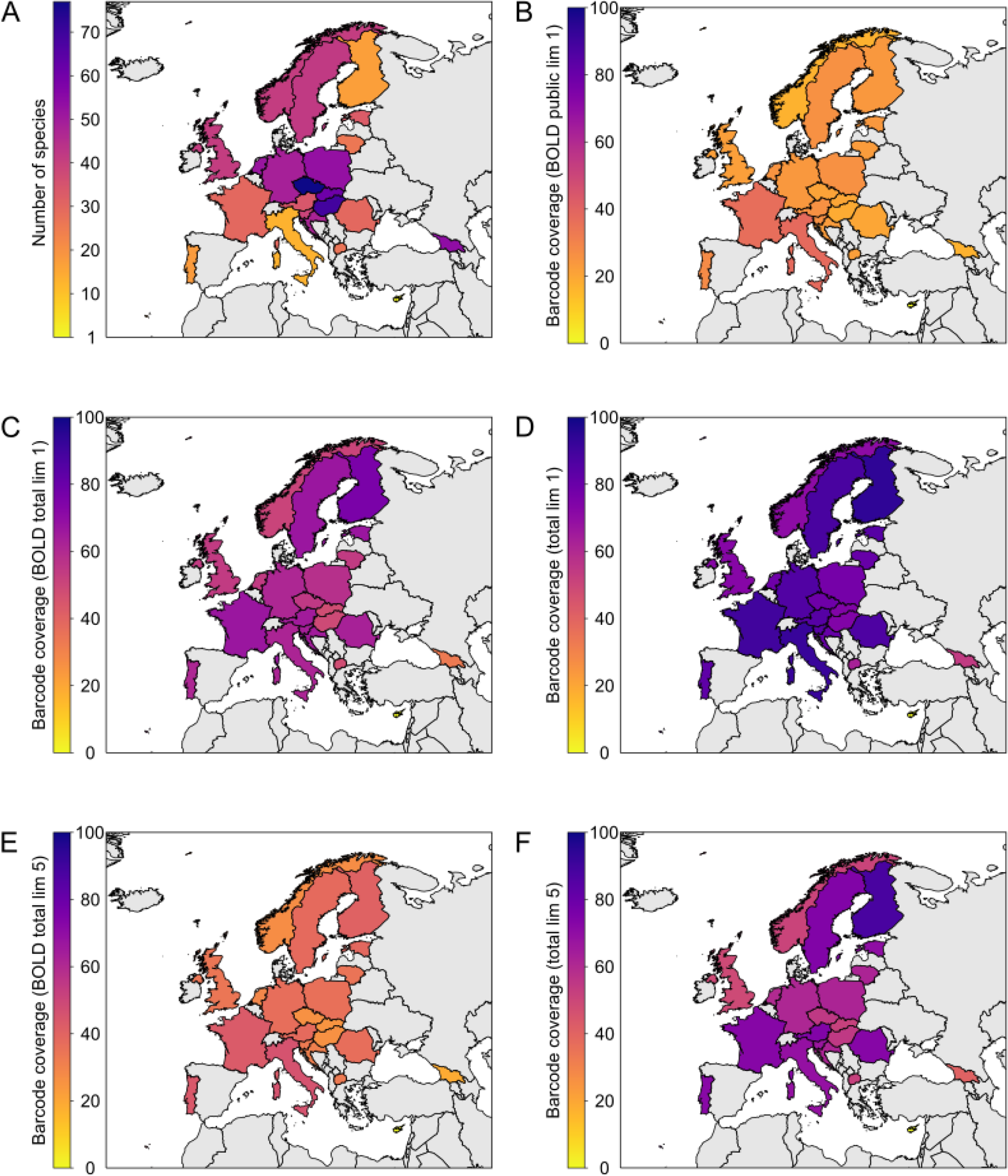
Barcode coverage maps of Mollusca. (A) Number of monitored species per country. (B) - (F) Barcode coverage per country for different datasets (BOLD public, BOLD total and total) and thresholds (lim 1 = minimum one record; lim 5 = minimum five records).

A high proportion of the missing barcodes was found for species that are used in freshwater monitoring in a single country (41 species, Fig. 9D). Only five of the 35 species surveyed in at least ten countries had no barcodes available. However, a comparatively high number of widely distributed and as such often listed freshwater molluscs (at least on 5 lists) do not yet have barcode data (23%). The barcode coverage per country was relatively evenly distributed, with an average coverage of 23% (min: 0% - Cyprus, max: 38% - Italy) when public barcodes in BOLD were considered, 56% (min: 0% - Cyprus, max: 76% - Finland) when public and private data on BOLD were used and 76% (min: 0% - Cyprus, max: 94% - Finland) for the full BOLD and GenBank datasets (Fig. 13).

#### 3.4.6 Platyhelminthes

Overall, 61 freshwater flatworm species are used for monitoring in 16 countries. The number of species monitored per country ranged from one (Estonia) to 39 (Czech Republic). While most species are observed in only a few countries, there are nine species on at least ten national checklists, with *Dendrocoelum lacteum* being the most common (14 countries; Supplement 2). The barcode coverage of freshwater Platyhelminthes was very low (4.9%, Fig. 8) as only three species had sequences deposited in examined databases. Of these, two species (*Dugesia cretica* and *Girardia tigrina*) had only one COI-sequence mined from GenBank, while 51 private barcodes were available for *Dugesia gonocephala*.

#### 3.4.7 Nematodes

Nematodes can be ascribed to both meio- and macroinvertebrate fauna depending on size of the respective freshwater forms. Only ten of the national checklists include (assumed) free-living and semi-parasitic forms, mostly on a coarse taxonomic level. The lists contained one taxonomically wrong classification (*Gordius aquaticus* as Nematoda instead Nematomorpha), one fish parasite, semi-parasitic forms of the family Mermithidae, and one higher order which is taxonomically no longer in use (Secernentea). Only the Romanian list contain two relevant nematode species (*Dorylaimus stagnalis* and *Tobrilus gracilis*). These are common in freshwater, and both are represented with barcodes in BOLD.

### 3.5 Freshwater fish

As of 1^st^ February 2018, the target list for European freshwater fishes contained 627 species including 18 extinct and 3 ‘extinct in the wild’ species. After the first BOLD checklist query against all available data, 110 of the 627 species were listed as in need of specimens, i. e. completely lacking DNA barcode references in BOLD (coverage: 82.5%, Fig. 14A). When setting the threshold for minimum number of DNA barcodes available to five, 212 species did not have any or fewer than five barcodes deposited in the database (Fig. 14B). After manually checking the resulting gap list and taking into account real synonyms and different taxonomic concepts such as generic assignments (e.g., *Iberocypris* vs. *Squalius*, *Orsinigobius* vs. *Knipowitschia*, only 60 extant species (plus 16 extinct) were not represented with DNA barcodes (coverage: 90.4% or 87.9% including extinct species). Only three species listed in BOLD had records that did not fulfil the formal requirements for DNA barcode status.

**Figure 14.**
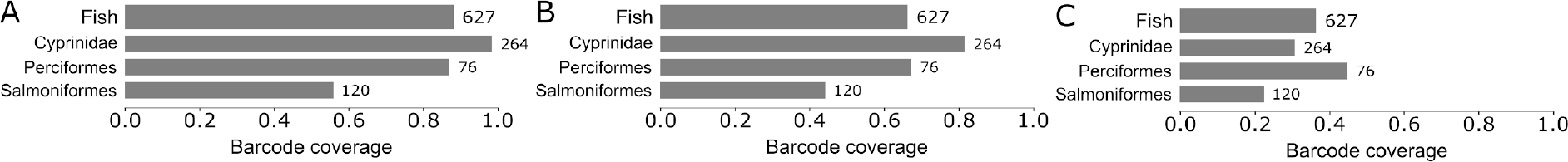
Barcode coverage for freshwater fish. A) A minimum of one DNA barcode, B) ≥5 DNA barcodes or C) one 12S sequence per species. Numbers on bars refer to the number of species in checklist. Eighteen extinct and 3 extinct in the wild species are included.

In general, the DNA barcode (COI) coverage for extant species is very good in most countries (100% coverage in 16 countries) and only a few species are missing reference records from certain regions (Fig. 15A, C). In the Scandinavia and the UK, a number of chars (*Salvelinus* spp.), trouts (*Salmo* spp.) and whitefishes (*Coregonus* spp.) are not yet represented in the databases. While for Austria, Germany and Switzerland, a smaller number of whitefishes (<10) are still missing in the DNA barcode reference libraries. Concerning extinct and extinct in the wild species (Fig. 15B), only a few species are missing, the highest number of them (6) reported from Switzerland.

**Figure 15.**
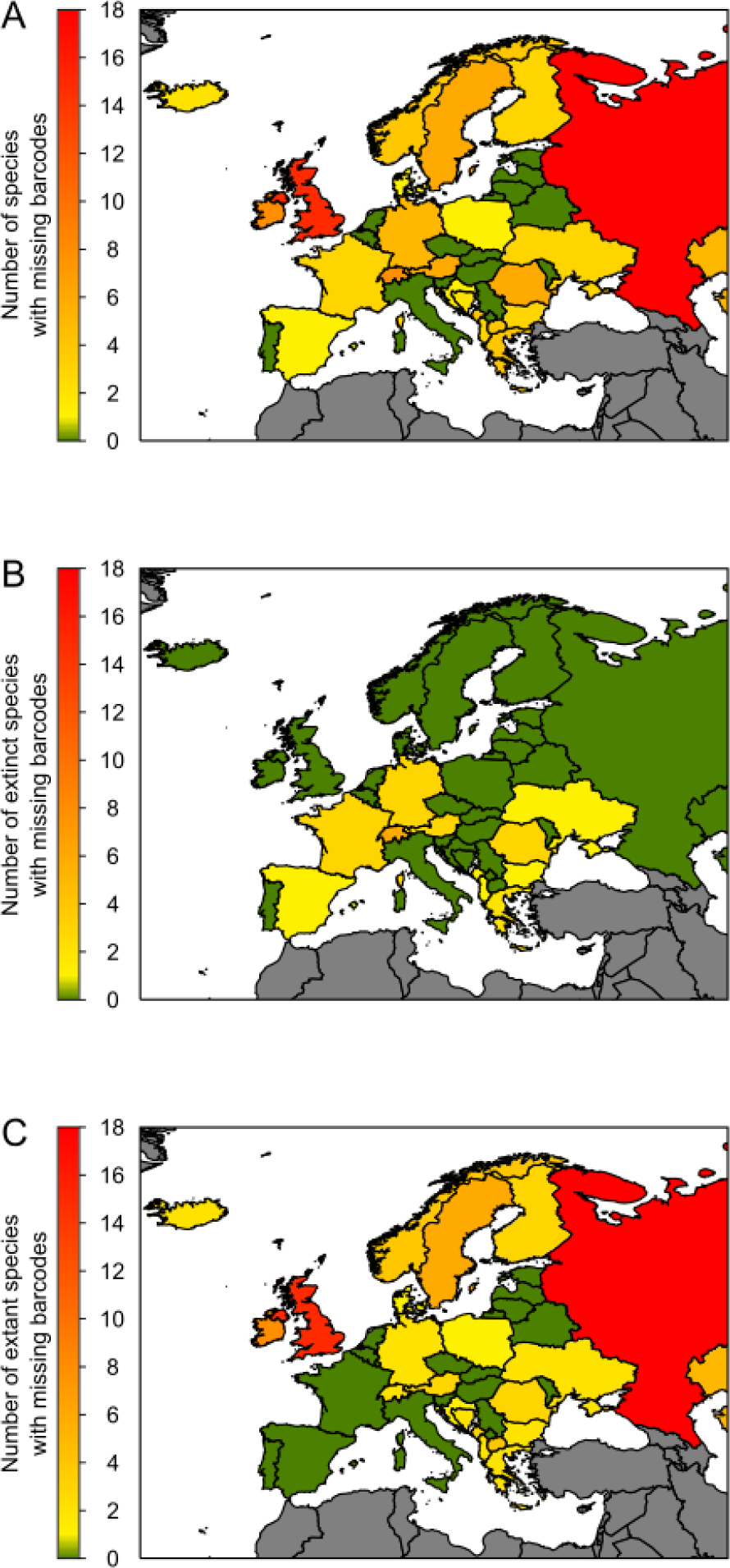
Missing barcodes for freshwater fish species. (A) Number of all species with missing barcodes per country. (B) Number of extinct species with missing barcodes per country. (C) Number of extant species with missing barcodes per country.

### 3.6 Reverse taxonomy

Documented use of reverse taxonomy was observed in all groups of freshwater macroinvertebrates where public data was available, except for Neuroptera (Fig. 16, Supplement 3). The proportion of identified sequences originating from reverse taxonomy compared to all available barcodes ranged from 1% (Crustacea, Ephemeroptera, Hemiptera, Lepidoptera and Odonata) to 20% (Coleoptera) and 59% (Diptera). Since these values rely on the cumulative number of BOLD-public, BOLD-private and GenBank data, and since the use of reverse taxonomy is know only from public sequences in BOLD, the calculated proportions can be underestimations. For instance, when only public data in BOLD is considered, reverse taxonomy can be found in up to 61% (Annelida) and 82% (Diptera) of the deposited sequences. The fraction of species with barcodes originating from reverse taxonomy ranged from 3% (Arachnida, Coleoptera and Ephemeroptera) to 16% (Diptera) and 20% (Megaloptera). Although the proportion of species having reverse taxonomy of potentially strong influence was low for most taxonomic groups, it was comparatively high for Diptera (12%) and Megaloptera (20%).

**Figure 16.**
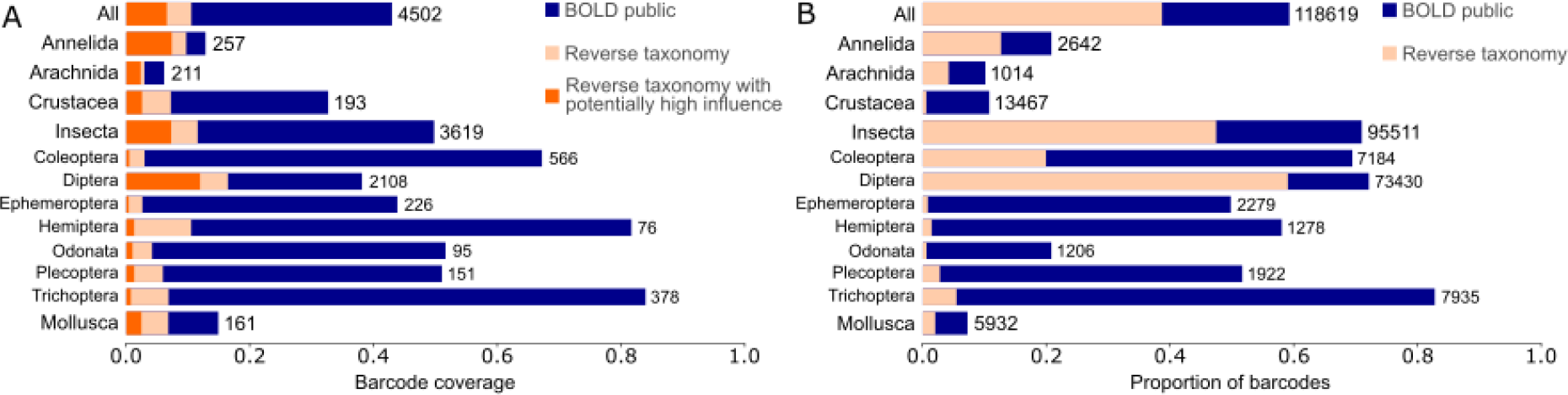
Overview of reverse taxonomy in freshwater macroinvertebrates. (A) The proportion of species with reverse taxonomy barcodes for the different taxonomic groups. (B) The proportion of barcodes originating from reverse taxonomy for the different taxonomic groups. Thick bars represent higher taxonomic ranks, thin bars represent insect orders. Numbers on bars refer to total number of (A) species in checklist or (B) total barcodes. Taxonomic groups with less than ten species or without public data are not indicated.

## 4. Discussion

### 3.1 National checklists

We collected lists of species that are used in the national implementations of the WFD and MSFD monitoring programs and water quality assessments in each target country. However, since countries have different strategies on how they report and comply with the regulations, the lists were very different in terms of taxonomic coverage and level of classification among nations. WFD monitoring requires the use of stressor-specific Multimetric Indices (MMIs, (Hering et al., 2006)), intercalibrated and validated at river basin level (Hering et al., 2010). Each country has developed their own set of MMIs, each consisting of biotic indices which are best suited to describe water quality status in their region (Birk et al., 2012a; European Commission, 2018). The taxonomic depth of data required for calculation of these indices is highly variable between countries. In cases where indices are dependent on species-specific traits, all species counts and complete species-level identification is required. Thus, the checklists of species from each country that we received and have used as basis for our gap-analysis can be grouped into four major types.

The first group contains ‘full national lists of species’. Such lists are typically generated from the Pan-European species lists, or compiled individually from literature. Some countries (e.g. Czech Republic) use these complete lists as basis for their WFD monitoring, even if many taxa are not regularly encountered. The second group includes lists from countries that use the national taxa lists as a basis, but narrow down the selection based on experience or challenges with identification to species level. In Hungary, for example, only species that were previously recorded during WFD monitoring are used. Other countries would limit the identification of selected groups to family- or genus-level, or completely discard semi-aquatic taxa or taxa that are non-aquatic but closely connected to aquatic environments (e.g. Carabidae, Chrysomelidae or Curculionidae beetles). These restrictions have been taken into account in index development. In the third group, it is common to regularly monitor the frequency/occurrence of certain ‘highly indicative’ species/taxa, and use only these species in the calculation of MMIs. Thus, a highly restricted ‘operational taxon lists’ for WFD monitoring is compiled. Such a list can be extensive or quite short, dependent on country. The fourth group includes lists that are exclusively based on family or genus-level identifications. The differences in the submitted taxa lists influence how the geographical coverage of DNA barcodes should be interpreted. It might be most obvious for the first group of lists, where a considerable number of taxa are not regularly encountered but still included in the barcode gap-analysis through the checklists from Sweden, Czech Republic and Slovakia (Figs 10-13).

### 4.2 Marine macroinvertebrates & fish

Based on the ERMS checklist, the gap in the reference library for marine species is relatively large (> 70%) for all analysed taxa, with the exception of fish (18%). The gap is much smaller when the AMBI checklist is used (little over 50%), but still comparatively high in global terms, particularly compared to libraries of freshwater taxa. The lower coverage of the ERMS list is somehow expected since this list contains six times more taxa than the AMBI list (18,451 versus 3,012, respectively). For the purpose of bioassessment under the WFD and MSFD, AMBI is a more relevant checklist since it describes the sensitivity of macrozoobenthic species to both anthropogenic and natural pressures, and it is currently used as a component of benthic invertebrates’ assessment in many EU member states within the four regional seas (Borja et al., 2007; Borja et al., 2009; European Commission, 2018).

The percentage of barcoded species greatly differed between lists and targeted taxonomic groups. Marine fish (only included in the ERMS list), were by far the best represented taxonomic group, with barcodes available for 82% of the nearly 1,500 species in the list. Partially due to the commercial importance of this group, marine fish have been the target of comprehensive DNA barcoding campaigns along multiple marine regions in Europe (e.g. (Costa et al., 2012; Keskİn and Atar, 2013; Knebelsberger et al., 2014b; Landi et al., 2014; Oliveira et al., 2016)). However, a fair proportion of the barcode records for marine fish may not have originated from specimens collected in European seas (Oliveira et al., 2016), since many species have considerably large distributions (e.g. (Ward et al., 2008)) including, for example, those occurring also in the northwest and south Atlantic Ocean.

For marine benthic macroinvertebrates in the AMBI list, the three most species-rich phyla; Annelida, Mollusca and Arthropoda (ca. 85% of the total species in the list), have moderate levels of completion (40% to 50%), while less represented groups such as Nemertea, Sipuncula and Echinodermata have completion levels of at least 65%. Within the ERMS list, the levels of completion were lower than those of the AMBI list, but followed similar trends of those reported for the AMBI list, with the exception of the nemerteans. The Annelida, Mollusca and Arthropoda, that accounted for ca. 77% of the species in the ERMS list, have fair levels of completion (20% to 30%) and lower than less diverse groups in the list, such as Echinodermata (35%) and Sipuncula (42%).

Our results suggest that many of the barcode studies focused on Annelida, Mollusca and Arthropoda, may have targeted particular species or groups at the order or family level (e.g. Crustacea (Costa et al., 2007; Raupach et al., 2015); Decapoda (Matzen da Silva et al., 2011a; Matzen da Silva et al., 2011b; Matzen da Silva et al., 2013); Amphipoda (Lobo et al., 2017); Gastropoda (Barco et al., 2013; Barco et al., 2016; Borges et al., 2016); Polychaeta (Lobo et al., 2016); Bivalvia (Barco et al., 2016). A closer look into particular taxonomic groups in our analysis supports this: for the order Decapoda, which comprises only 25 species in the AMBI list and 693 species in the ERMS list, 84% of the species are barcoded in the former, and ca. 50% in the latter. For a larger group such as the superorder Peracarida, which comprises 649 species in the AMBI list and 2,643 species in the ERMS list, the total number of barcoded species is much far from completion (45% and 24%, respectively).

In addition to the globally modest levels of completion for marine macroinvertebrates, the gap-analyses based on the AMBI checklist also reveals some insufficiencies of the available data, namely the presence of a sizeable proportion of private records, which are unavailable for full access in bioassessment studies employing DNA-based tools. For some groups, private records on BOLD were even higher than the public, such as for Sipuncula (25% *versus* 10%) and Annelida (20% *versus* 18%). An ISI Web of Science search, at the time of writing (30^th^ November 2018), with the search terms “barcoding” AND “marine” AND “the taxonomic group of interest” also supports the absence of published reference libraries for Sipuncula, or the low number of studies found for Annelida, compared to other above-mentioned groups (e.g. fish and Crustacea). Another aspect worth of consideration is the number of singletons in the reference libraries. Although the percentage of singletons is generally low, some taxa have a considerable proportion of single representatives per species. Whereas relatively low levels of barcode coverage for some of these groups clearly reflect fewer efforts to barcode those taxa, a considerable proportion of the gap must also be ascribed to failed DNA sequencing, due to either primer mismatch, sample contamination or PCR inhibitors. This is particularly obvious for the marine Annelida, for which COI sequencing success rates may be down to 40-50 % on average (Kongsrud et al., 2017). Barcoding of annelids has also revealed unexpected high levels of genetic diversity, prompting traditional species taxa to be torn apart (Nygren, 2014; Nygren et al., 2018). A relatively high proportion of private data may reflect that some species taxonomies are currently in a certain state of flux. [ref. to this insert:

By increasing the threshold of at least one barcode per species to five barcodes, the level of completion of both lists (i.e. ERMS and AMBI) fell to about half. For instance, the levels of completion remained acceptable only for fish and Decapoda, but for most groups these are greatly distant from what would be recommendable, in particular for Sipuncula, Nemertea, Cnidaria, Brachiopoda and Annelida. Ideally, reference libraries should have a fair and balanced representation of specimens across the geographic distribution for each species, to capture the range of intraspecific variation in the DNA barcodes in the best possible way. Such representation is also key for efficient quality assurance, quality control and validation of reference libraries, as discussed below.

Within the AMBI list, almost half of the species fall into the ecological group I, which are the “sensitive” species, and the remaining half is distributed among the other 5 ecological groups. However, the completion levels were higher for species from ecological groups III (56%) and V (52%) and lower for species that do not have any ecological group assigned (38%). Similar results were encountered when the first attempt of using a genetics based marine biotic index (gAMBI), with available GenBank sequences for AMBI species, has been performed (Aylagas et al., 2014). At the time, the authors concluded that the available genetic data was not sufficient or did not fulfil the requirements for a reliable AMBI calculation, that needs an even distribution of taxa across the disturbance gradient. On the other hand, when gAMBI values were calculated by using the most frequent species within each ecological group, the reliability of AMBI values increased significantly (Aylagas et al., 2014). Nevertheless, in the current study we have found a much higher completion level (e.g. 48% versus 14%), since numerous new records have been generated in the meantime and our gap-analyses also included BOLD data.

### 4.3 Diatoms

Several diatom studies have pointed out the barcode reference library as the Achilles heel of using metabarcoding of diatoms for environmental monitoring (Kermarrec et al., 2013; Rivera et al., 2018a; Rivera et al., 2018b; Vasselon et al., 2017). The barcode reference library must be as comprehensive as possible in order to assign a high proportion of environmental sequences to known taxa, and it requires regular expert curation in order to maintain quality. This is why experts from several countries joined efforts to curate a single reference library, Diat.barcode (formerly called R-Syst::diatom). Our results show that a large majority of the most common species (registered in the checklists of all countries) are present in this library, but that many rare species lack representation.

A comprehensive barcode reference library for diatoms is difficult to achieve for two reasons. Firstly, because more than 100,000 species are estimated to exist globally (Mann and Vanormelingen, 2013), many of which are undescribed. Registration of barcodes and metadata of all these species in the reference library will require an overweening effort. Thus, an effort should be focused on the most common, not yet barcoded species. Secondly, diatoms need to be isolated and cultured in order to obtain high quality, vouchered, barcode records. This work is tedious and often unsuccessful because many species are difficult or impossible to cultivate. As a remedy to this, an alternative method using high throughput sequencing of environmental samples was proposed by (Rimet et al., 2018b). By using this method routinely, we will be able to quickly complete the barcode reference library of the most common diatoms in the near future.

### 4.4 Vascular plants

For vascular plants the standard DNA barcode is the combination of two plastid loci, *rbcL* and *matK*. Logically, this simple fact doubles the effort needed when barcoding plants. Fortunately, some national campaigns of flora barcoding have been developed in the last decade (e.g. http://botany.si.edu/projects/dnabarcode/index.htm; https://botanicgarden.wales/science/saving-plants-and-fungi/dna-barcoding/; https://www.rbge.org.uk/science-and-conservation/scientific-and-technical-services/dna-barcoding/dna-barcoding-britains-liverworts/) and the vascular plant species used for water quality assessments are well represented in the public databases (BOLD, GenBank), with barcodes registered for more than 83% of the species. The gap-analysis tool on BOLD that was used here does not require both loci to be barcoded for plants, as it reports the percentage of barcoded species regardless of whether sequences exist for both loci or only one. Only a manual check on the public data in BOLD could overcome this problem, whereas no information can be obtained for private data about the barcoding marker used. With a total of 515 barcoded species, the locus *rbcL* is better represented than *matK* (449 species). Amplification and sequencing of the *matK* barcoding region is widely known to be difficult due to high sequence variability in the primer binding sites (Hollingsworth et al., 2011). Considerable efforts have been made for developing efficient primers across multiple angiosperm families, as reflected in the recent study published by Heckenhauer and colleagues (Heckenhauer et al., 2016).

In order to have a complete evaluation of the state of DNA barcode data for the macrophytes, analyses should also be performed for charophytes and bryophytes. One should, however, be aware that the situation is far from simple. A universal DNA barcode has yet to be identified for bryophytes, for which commonly used markers have low amplification-sequencing success or lack of resolving power at the species level (Hassel et al., 2013). As for the charophytes, species morphological delineation might be complicated given the plasticity of the discriminatory characters. Recent studies based on DNA barcode analyses showed that differentiation of closely related *Chara* species is not always possible and questioned the relevance of certain morphological traits in the species differentiation (Schneider et al., 2015), by highlighting an incomplete process of speciation (Nowak et al., 2016).

### 4.5 Freshwater macroinvertebrates

Macroinvertebrates are central BQEs in freshwater biomonitoring programs. Our barcode gap-analyses of the BOLD reference library, including data mined from GenBank, show that while there is comparatively few species missing sequences in some insect orders (e.g. Hemiptera, Odonata and Trichoptera), other taxonomic groups lack barcodes for a majority of the regularly monitored species (e.g. Platyhelminthes). Diptera, the most species rich group used in biomonitoring in Europe, had a fairly low coverage with only about 60% of the species represented in the reference libraries. This result is similar to what was recorded in a gap-analysis of the North American freshwater invertebrates (Curry et al., 2018), although their analysis was done on genus-level. In a barcode-gap analysis of the Great Lakes fauna Trebitz et al. (2015) found that rotifers, annelids and mites had particularly low coverage, while about 70% of all insect species were represented by barcodes in BOLD. While these numbers might have changed by now, it is interesting to see that the coverage of mites and annelids appears better in Europe, while insects are slightly better covered for the Great Lakes. Generally, in our results, there is a pronounced increase in taxonomic coverage when private data in BOLD and GenBank data are included. This is particularly obvious for Annelida, Arachnida, Crustacea and Mollusca (Fig. 8). It should also be noted that while species-level coverage is low for some groups, coverage often increases at higher taxonomic ranks. This is of relevance, as some taxonomic groups are only reported at the genus, family or even order level by several countries. Below we discuss some of the characteristics observed for each major taxonomic group.

#### 4.5.1 Insects

Insects are among the or even the most important and most often monitored organisms in freshwater assessments. This is reflected by both a high number of countries monitoring insects and a high number of monitored species in national monitoring checklists. However, the taxonomic level applied as well as the number of monitored taxa differs vastly among countries. The differences typically reflect the national monitoring programs (Birk et al., 2012a; Kelly et al., 2015) and hinder a direct comparison of countries in many cases (requiring sophisticated intercalibrations) and also affect the overall gap-analysis result. Almost two-thirds (65.7%) of the monitored insect species are barcoded. When looking at the taxa with the lowest barcode coverage, it becomes apparent that most of the missing species (70%) belong to Diptera, of which 72.9% are exclusively monitored in a single country (Czech Republic). Excluding only these missing Diptera species from the gap-analysis increases the overall coverage from 65.7% to 79.7% of the species, rendering the observed gap in the barcode coverage partly a problem resulting from one excessively long national checklist. This is further supported by the fact that otherwise, on average, 88.5% of the monitored species across all other surveyed countries have sequences in the reference libraries. However, similar to the observations made by Trebitz et al. (2015) for the Great Lakes fauna, the barcode coverage is significantly reduced when considering species that are represented by at least five barcodes. Moreover, since regional coverage in barcode reference libraries is important to account for the genetic diversity that is correlated with geographic distance (Bergsten et al., 2012), geographic coverage maps (Fig. 10, Suppl. Figs 3-9) can be useful to identify priority areas when filling gaps in the barcode library. For some countries (e.g. Georgia), the low coverage of barcoded species can be explained by many unique species in their national checklist. In such cases, regional representation in the barcode library is crucial for implementation of DNA barcoding in freshwater biomonitoring.

One obvious discrepancy was observed for the common mayfly *Ephemera danica.* While this species is one of the two most monitored species, there are only 15 available barcodes in BOLD despite there being 151 registered records. The low sequencing success of this species can be explained by suboptimal lab protocols (e.g. primer cocktails), and better representation of this commonly monitored species on BOLD could probably be achieved through protocol optimization. In conclusion, even if gaps still need to be closed, the reference databases for insects in Europe are well developed making this group already qualified for monitoring through DNA metabarcoding in several countries (e.g. (Morinière et al., 2017)).

#### 4.5.2 Arachnids

Aquatic arachnids are not commonly monitored in Europe, at least not for the WFD. The most species-rich group, water mites, are well suited for monitoring environmental change of many habitats (Cantonati et al., 2006; Gerecke and Lehmann, 2005). Species-level identification using molecular tools will make information from this group more readily available in the near future. Currently, most of the barcode data on water mites in BOLD are private, but the coverage is relatively high (Fig. 9B) thanks to efforts in the Netherlands and Norway (pers. obs.). Barcode data has revealed taxonomic challenges in water mites, as revealed by the 18 specimens of *Lebertia porosa* from Norway that comprise 7 BINs (Stur, 2017), and show a mean intraspecific p-distance of 11.7% (max 18.5%). Knowing that this species has currently 27 taxonomic synonyms, it will need some efforts to disentangle the names potentially associated with each genetic cluster. For the species *Hygrobates fluviatilis* a similar situation was solved with the help of DNA barcodes (Pešić et al., 2017). It is notable that the divergence of lineages with potentially different environmental preferences within the *H. fluviatilis* complex would not have been easily discovered without the comparison of sequence data in a barcode library.

#### 4.5.3 Crustaceans

Crustaceans, predominantly malacostracans, are quite commonly monitored in European countries. However, the level of their taxonomic identification varies a lot from country to country, and depends on the crustacean group considered. The species are generally well covered in BOLD, however, almost half of them are represented by private barcodes only. As such, they potentially form parts of large datasets deposited in BOLD in result of ongoing studies, which eventually will be published soon. Still a comprehensive DNA barcode library for European freshwater crustaceans, such as the one published for marine crustaceans (Raupach et al., 2015), is far from completion. Yet, there are numerous recent publications providing a wealth of DNA barcode sequences as a side effect of phylogeographic or taxonomic studies, revealing the presence of high cryptic species diversity in numerous morphospecies (e.g. (Christodoulou et al., 2012; Mamos et al., 2016). For groups such as amphipods, publication of barcodes along with descriptions of new species and cryptic lineages has become almost a rule (e.g. (Rudolph et al., 2018). Thus the prognosis for further extending the reference libraries in a foreseeable future is positive.

#### 4.5.4 Annelids

Despite the fact that numerous species of annelids are monitored in European countries, they are poorly covered in BOLD and most of the barcodes are kept private. A substantial share of barcode sequences mined from GenBank only. It seems that so far, there is no general habit of using BOLD as a repository for sequence data, even though COI barcodes were proven useful for identification of pseudo-cryptic and cryptic species of medicinal leeches almost a decade ago (Phillips and Siddall, 2009; Siddall Mark et al., 2007). Soon thereafter, an incongruence between morphological and molecular species boundaries was proven for *Erpobdella* leeches (Koperski et al., 2011). More recent studies revealed substantial cryptic diversity within several genera and species of freshwater oligochaetes (e.g. (Liu et al., 2017; Martin et al., 2018; Martinsson et al., 2013; Martinsson and Erseus, 2018). Thus, it appears that DNA barcoding would be immensely beneficial for identification of annelids in biomonitoring.

#### 4.5.5 Molluscs

A remarkable finding for freshwater molluscs was their comparatively high number of DNA barcodes deposited in GenBank, and not in BOLD. This pattern can be interpreted in terms of early initiated molecular taxonomic endeavours in the pre-BOLD era, or by a community-behaviour of submitting sequences to GenBank rather than to BOLD (e.g. (Benke et al., 2011; Prie et al., 2012). When doing so, relevant metadata might be omitted or not immediately linked to the barcode. Thus, direct BOLD submissions are highly encouraged. Furthermore, a considerable proportion of frequently listed, and presumably widely distributed species do not yet have any barcode data available. This lack of data might be even more pronounced, as several integrative taxonomic studies on freshwater molluscs indicate that widely distributed morphospecies often comprise complexes of distinctly defined genetic lineages (cryptic species). A good example is *Ancylus fluviatilis*, the most often listed freshwater mollusc in our dataset, which actually constitutes a complex of at least six cryptic species (Albrecht et al., 2006; Pfenninger et al., 2003; Weiss et al., 2018).

#### 4.5.6 Platyhelminthes and Nematoda

Both flatworms and nematodes are diverse and of indicative value. While some countries do register Platyhelminthes in existing surveys, Nematoda are generally neglected. For nematodes in the Palaearctic, 1580 species should be relevant for the WFD (Eisendle et al. 2017). Thus, a barcode library of freshwater nematodes can have a potentially large impact on the use of this organism group in future biomonitoring.

### 4.6 Freshwater fish

With about 88% coverage with at least one DNA barcode in BOLD, European freshwater fishes are well represented and the species being reliably identifiable based on COI in real world applications. While 47 species have only one specimen with DNA barcode deposited in BOLD, a large proportion (two thirds) is available with at least five individuals. The coverage with 12S data is much lower and only a third of the species was found to be available in public databases.

About three-fourth of all European freshwater fish species fall into the three higher taxa presented in more detail (Perciformes, Cyprinidae and Salmoniformes), which contain commercially important game and food species (perch, pike-perch, carp, bream, roach, trout, whitefishes and chars). The largest and most widespread family is Cyprinidae, which has its (extant) species completely covered by DNA barcodes in BOLD, with only five species missing - all of which being regarded or listed by IUCN as extinct (*Alburnus danubicus*, *Chondrostoma scodrense*, *Iberocypris palaciosi*, *Pelasgus epiroticus*, *Romanogobio antipai*). Especially given the potential of molecular identification and detection tools for non-invasive and highly sensitive approaches to assess a species’ presence or even abundance (e.g. (Ushio et al., 2018), we argue that it is also important to cover those species in the databases, which are thought to be extinct. Among the ten completely missing perciform species are four tad-pole gobies (*Benthophilus* spp.), two sculpins (*Cottus* spp.) and two dwarf gobies (*Knipowitschia* spp.) with predominantly Eastern European and putative Caspian basin distributions, areas which are generally less well studied and explored from an ichthyologist’s perspective. An exemption is the elusive *Zingel balcanicus* from Macedonia and Greece (protected through Annex II of the European Union Habitats Directive 92/43/EEC), which has been re-discovered and anatomically analysed recently (Arsovska et al., 2014), but as no DNA-material has been secured cannot be assessed via molecular tools at the moment.

The largest gaps in the reference database pertain to the salmoniform group with chars (*Salvelinus* spp. ‑ 19 species), trouts (*Salmo* spp. ‑ 9 species) and whitefishes (*Coregonus* spp. ‑ 16 species). While these groups contain many commercially exploited species, they are known to be notoriously difficult to identify based on general morphology (Kottelat and Freyhof, 2007), but also applying standard DNA barcoding routines (Geiger et al., 2014; Knebelsberger et al., 2014a). This is most likely due to the presence of post-glacially evolved species flocks, which are only little differentiated genetically ‑ at least judging from the groups that have been studied so far (Dierking et al., 2014; Hudson et al., 2011; Vonlanthen et al., 2012). From a geographic point of view, most missing species occur in Scandinavia and UK (chars), the Alp region (whitefishes), and the Eastern Mediterranean (trout).

### 4.7 Quality measures for DNA barcode reference libraries

In a barcoder’s perfect world, all species on Earth would be identifiable based on their DNA barcodes. However, this ideal conception is hampered by several biological and human-made phenomena. For example, time since speciation might be rather short, and the universal marker considered not diverse (i.e. not informative) enough to resolve this speciation event (e.g. (Weiss et al., 2018)). Additionally, gene flow might be still possible, even between less closely related species, leading to the (unidirectional) introgression of genomes, and hence to the (partial) intermixture of barcodes (e.g. (Weigand et al., 2017)). Besides these natural processes complicating the diagnostic utility of DNA barcodes, human-made artefacts during reference library development directly affect the reliability of DNA barcoding to correctly identify specimens to species. This includes identification errors, sequence contamination, incomplete reference data or inadequate data management. It was thus not surprising that subsequent to the proposal of the term “DNA barcode” (Floyd et al., 2002; Hebert et al., 2003a; Hebert et al., 2003b), special emphasis was laid on formal standardisation guidelines for DNA barcodes in the context of reference library development (see e.g. (Ratnasingham and Hebert, 2007; Walters and Hanner, 2006). Those include the criteria that any ‘formal’ barcode sequence: a) derives from an accepted gene region, b) meets certain sequence quality standards (e.g. demonstrating at least 75% of contiguous high quality bases or less than 1% Ns and being associated with trace files and primer information), c) is linked to a voucher specimen in a major collection, and d) ideally but not always mandatorily possesses further collection and identification details (i.e. georeference data, name of collector and identifier). Since then several biomonitoring and assessment applications have moved from classical single specimen identifications to highly parallelized characterisations of communities via DNA metabarcoding (Leese et al., 2018). Given the often overwhelming quantity of ‘big biodiversity data’ and automated pipelines in those HTS approaches, data quality aspects of DNA barcode references gain an even higher relevance. Thus, some research communities, such as European diatom experts have worked with the European Standardization Committee to publish a methodology as a first step towards standardization of reference barcode libraries for diatoms (CEN, 2018).

In principle, two quality components can be distinguished: Quality assurance (QA) is process-orientated, providing and maintaining quality standards for DNA barcodes and reference libraries. Quality control (QC), on the other hand, is user-orientated, enabling the cross-validation of taxonomic assignments or flagging of doubtful barcodes. More generally speaking, QA and QC measures can be seen as internal (or preventive) and external (or reactive) curation of reference libraries, respectively (Fig. 17). The implementation of QA measures during reference library development is the first important step for a sustainable data quality management. Linked to a valid taxonomy, formally-correct barcode sequences are deposited in line with (digital) voucher specimens and extensive metadata information. The taxonomic backbone should be regularly updated with modifications being visible to the users. An open access and fully transparent reference library allowing for versioning of barcode collections and the possibility to track taxonomic changes can be seen as the gold standard here. Simultaneously, this will allow a more sophisticated QC by the barcoding community. Library entries can be flagged for contamination and the most recent taxonomic changes (i.e. newly described species, integrative revisions) incorporated into the reference library taxonomic backbone more easily. A library which communicates with other ecological or geographic datasets and which provides access to the full data lifecycle from deposition to publication of data will further smoothen the integrative utilisation of barcode datasets. The generation of custom reference libraries and their annotation with digital object identifiers (DOI) finally can account for transparency and the specific demands of the users.

**Figure 17.**
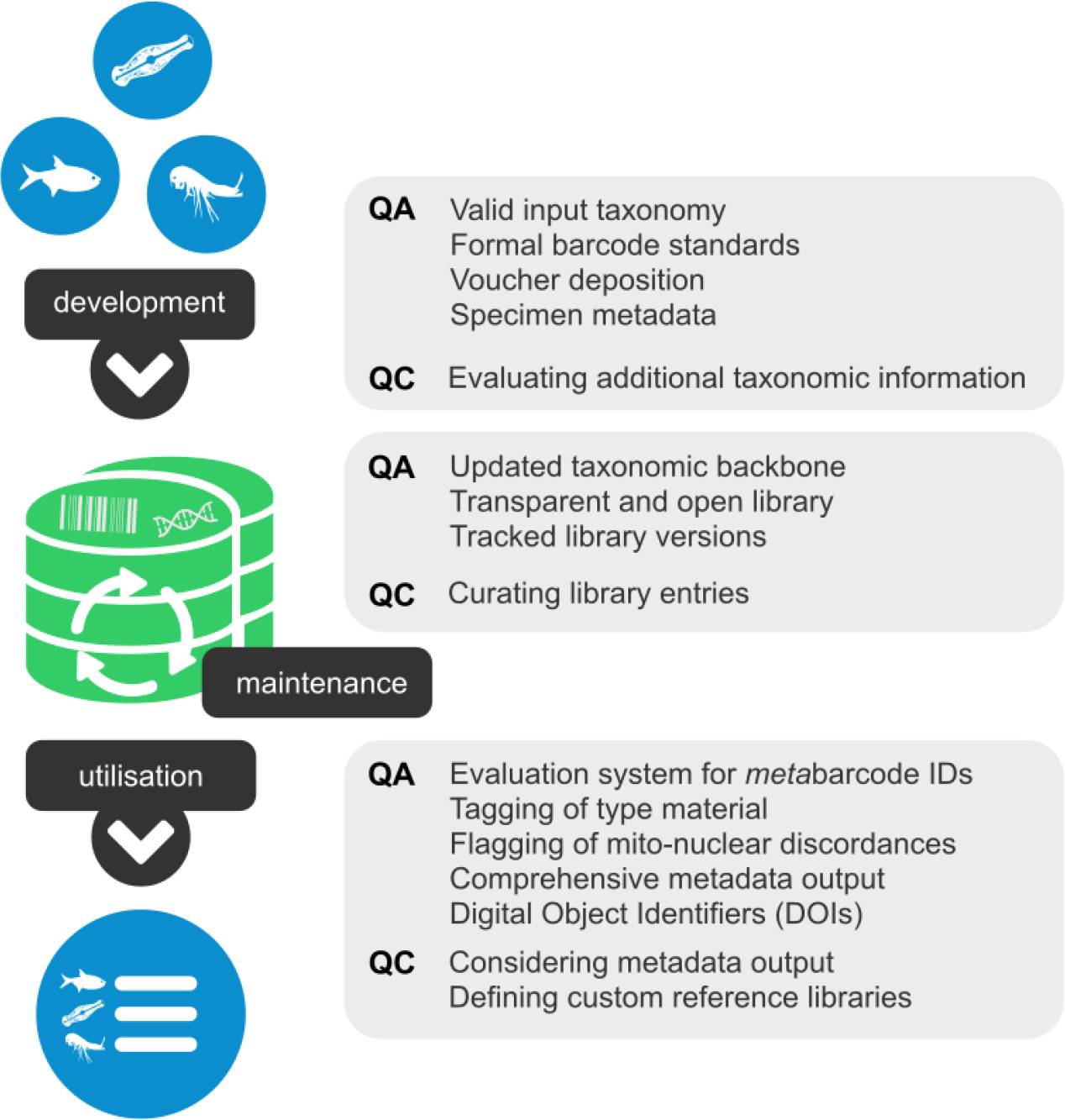
An overview of the reference library steps ‘development’, ‘maintenance’ and ‘utilisation’, their quality assurance (QA) and quality control (QC) measures.

Although a variety of QA/QC measures are implemented at the stages of reference library development and maintenance, improvements are possible for the QA/QC components during reference library utilisation. This holds especially true for complex metabarcoding studies based on multiplexed HTS data. In most of those cases, taxonomic identifications are achieved by semi-automatically comparing the clustered or individual metabarcodes with a reference library and applying flexible similarity thresholds. The sequence is thus linked to a Linnaean name, e.g. by a 2% similarity threshold for species-level identification of a molecular operational taxonomic unit (MOTU). By doing so, only the availability and match of barcodes are considered, neglecting any additional metadata. Yet, knowledge about the number of barcoded specimens per species, their morphological identifiers and the distribution area covered, are likewise valuable information and should be available for direct QC. Special cases of mito-nuclear discordance, the number of already known MOTUs for a given Linnaean species name and ‘extraordinary’ barcodes such as those originating from type specimens should be additionally highlighted in the output results. All this combined information could be used to establish an evaluation system for metabarcode identifications, sorting taxonomic results by their plausibility and hence establishing further QA for reference library identification performance.

The ultimate reference library goal is to link a DNA barcode to a voucher specimen, accompanying metadata and its Linnaean name. However, more and more frequently, a reverse taxonomic approach is applied for the generation and deposition of reference barcodes, e.g. the ‘reverse BIN taxonomy’ in BOLD. During this process, a sequence with a taxonomic annotation above the species-level (e.g. family or genus) is included in the database and identified by the already available barcodes. For instance, a chironomid specimen (BOLD sequence page: GMGMC1513-14) of a vouchered collection in BOLD bears the species name *Polypedilum convictum* including a specimen identifier in its metadata. Only when accessing the internal specimen page (BOLD: BIOUG16529-F11) the identification method information “BIN Taxonomy Match” is given, however, without presenting the original morphological identification level. Strictly speaking, species identification through DNA barcoding has generated this ‘reference’, and not expert identification by morphology. Subsequently, this sequence is considered on species-level in the database and a more precise initial morphological identification is pretended. At present, reverse BIN taxonomy sequences (see Supplement 3) a) can be found in up to 16% and 20% of the monitored species of a taxonomic group (Diptera and Megaloptera, respectively), b) represent up to 61% and 82% of a higher taxon’s public barcodes (Annelida and Diptera) and up to 20% to 59% of all barcodes (Coleoptera and Diptera), c) can be found in species with only few public barcodes (e.g. three out of five *Crangonyx pseudogracilis* sequences are reverse BIN taxonomy sequences) and d) represent more than half (e.g. 38/75 for *Mytilus edulis*) or all (e.g. 35/35 for *Lumbricillus rivalis*) public sequences of a species. As such, wrong species-level DNA barcodes are potentially introduced, with often incorrect metadata for ‘specimen identification’ going along with them. They must be seen as a geographic reference for a MOTU rather than as a reliable taxonomic reference. The ‘reverse BIN taxonomy’ practice will also bias the evaluation system for the interpretation of metabarcode identification results.

An auditing and annotation system for reference libraries of DNA barcodes has been originally proposed by Costa et al. (Costa et al., 2012), and later updated by Oliveira et al. (2016) to accommodate the BIN system. The application of this QC system was particularly adequate for reference libraries of marine fishes (Cariani et al., 2017; Oliveira et al., 2016), but it has been equally applied to other taxa, such as Gastropoda (Borges et al., 2016) and Polychaeta (Lobo et al., 2016). Essentially, this system lies in the verification of the concordance between morphology-based identifications and BIN-based sequence clusters – within a given reference library (e.g. fishes of Europe) – and the subsequent annotation of each species with one to five available grades, i.e. ranging from maximum concordance (grade A) to complete discordance (grade E). Annotated grades are ought to be regularly reviewed and updated as required. Rather than requiring decisions about the taxonomic status and validity of a given species, this procedure simply considers the annotation of the level of congruency between morphological and molecular data. The auditor only needs to make decisions on the grade of congruency to apply.

The auditing system of Costa et al. (Costa et al., 2012) differs in a number of ways from the “BIN discordance report” tool implemented in BOLD, which only flags BINs that include records with more than one taxon name, but does not point out cases of the same species occurring in multiple BINs (note that BIN discordance reports of all data on BOLD only is available in BOLD v3). Also, because the BIN discordance report is an automated computer-based procedure, it does not distinguish true discordance from misspelled species names, synonyms, or patent cases of discordance resulting from cross-contamination or mislabelling of samples (e.g. (Knebelsberger et al., 2014b)). Hence, as a result of the auditing and annotation framework, end-users will have an indication of the reliability and accuracy of a given species match, and will be immediately alerted for records with insufficient data, or uncertain or misleading matches.

## 5. Conclusions and Recommendations

For marine macroinvertebrates, future efforts should focus initially on filling the gaps of the AMBI checklist, especially those more dominant in the datasets, which greatly influence the AMBI result (Aylagas et al., 2014), while keeping the long-term goal of completing the ERMS checklist. For freshwater macroinvertebrates, species-groups that are widely used in WFD monitoring such as Annelida, Crustacea, Insecta and Mollusca should be prioritized. For marine groups, gaps should be filled first to maximize phylogenetic representativeness, thereby yielding to the collection of reference barcodes of representative species from missing orders, then missing families, and so forth down to genera. This strategy aims to provide, at the very least, an interim proximate taxonomic assignment for metabarcoding reads lacking species level matches. However, most of the work has still to be done at the species level, because within the same genus, there are species belonging to different ecological groups, and thus the identification at species level is mandatory for reliable EQS and environmental status assessment. Hence, subsequent efforts should address species level completion, focusing on the taxonomic groups with greater gaps, as well as on the taxa used in AMBI’s ecological categories. The increase in the number of DNA barcodes for less barcoded species must also be pursued, since most of the taxonomic groups have less than 5 barcodes/species in the reference libraries. Attempts to include representative specimens across the geographic distribution range shall be made for missing species in the reference libraries. Particular care must be taken regarding the QA/QC of the reference barcode records to be produced, as failure to do so will limit their application, render them useless, or even introduce wrong outcomes. Moreover, as new HTS techniques are developed to obtain full-length reference barcodes from old type material (Prosser et al., 2016), this strategy should be used to resolve the taxonomy and names of key taxa used in biomonitoring.

## 6. Acknowledgements

This paper is a deliverable of the European Cooperation in Science and Technology (COST) Action DNAqua-Net (CA15219) Working Group 1, lead by Torbjørn Ekrem and Fedor Čiampor. Thanks to the University of Minho and University of Pécs for hosting workshops and working group meetings. We also thank staff at National Environment Agencies and others that provided national checklists of taxa used in biomonitoring, and otherwise assisted with checklist proof-reading: Jarmila Makovinská and Emília Mišíková Elexová (Slovakia); Steinar Sandøy and Dag Rosland (Norway); Mišel Jelič (Croatia); Marlen Vasquez (Cyprus); Adam Petrusek (Czech Republic); Kristel Panksep (Estonia); Panagiotis Kaspiditis (Greece); Matteo Montagna (Italy); Marija Katarzyte (Lithuania); Ana Rotter (Slovenia); Rosa Trabajo (Spain); Florian Altermatt (Switzerland); Kristian Meissner (Finland), Rigers Bakiu (Albania), Valentina Stamenkovic and Jelena Hinic (Macedonia); Patricia Mergen (Belgium); Gael Denys & the French Biodiversity Agency (France); Mary Kelly-Quinn (Ireland); Cesare Mario Puzzi (Italy); Carole Fitzpatrick (United Kingdom); Simon Vitecek (Austria); Ana Filipa Filipe (Portugal); Peter Anton Stæhr & Anne Winding (Denmark); Michael Monaghan (Germany); Alain Dohet, Lionel L’Hoste, Nora Welschbillig & Luc Ector (Luxembourg), Lujza Keresztes, (Romania). The authors also want to thank Dirk Steinke for providing the original European ERMS list for marine taxa and Florian Malard for comments on the manuscript. The preparation of the AMBI checklist was carried out in the scope of a Short-term Scientific Mission (ECOST-STSM-CA15219-150217-082111) granted to Sofia Duarte visiting AZTI Tecnalia, Spain.

## 7. Supplementary files

Supplement 1. Overview of obtained checklists of taxa used in national biomonitoring of aquatic ecosystems.

Supplement 2. Raw data from gap-analysis of all taxonomic groups showing: Species, Countries, Barcodes BOLD (public, private), GenBank records.

Supplement 3. Reverse BIN taxonomy statistics in BOLD.

## Author Contributions

All authors contributed to conceptualization and review of the manuscript. HW, AJB, FC, FOC, ZC, SD, MFG, MG, FR, BRu, MS, NS, AMW, EW, SAW, ZC-Z, SF, UE, JF, PG, WG, AH, BBH, BJ, JM, TM, GP, MAP, BWP, BRi, MALT, GV and TE provided and validated data. HW analysed data. TE, HW, FOC, MFG, AMW, SD, MG, FR, FC, MS, SAW, ZC, AJB and BRu drafted the manuscript. TE and FC administered the project and coordinated workshops.

## Footnotes

1 Oslo/Paris Convention on the Protection of the Marine Environment of the North-East Atlantic https://www.ospar.org/convention

2 Helsinki Convention on the Protection of the Marine Environment of the Baltic Sea Area http://www.helcom.fi/

3 United Nations Environment Programme - Mediterranean Action Plan to the Barcelona Convention http://web.unep.org/unepmap/

4 The Convention on the Protection of the Black Sea Against Pollution http://www.blacksea-commission.org/_convention.asp

