## Supplementary figures and images for "DNA barcode reference libraries for the monitoring of aquatic biota in Europe: Gap-analysis and recommendations for future work"

### Supplement Fig. 1

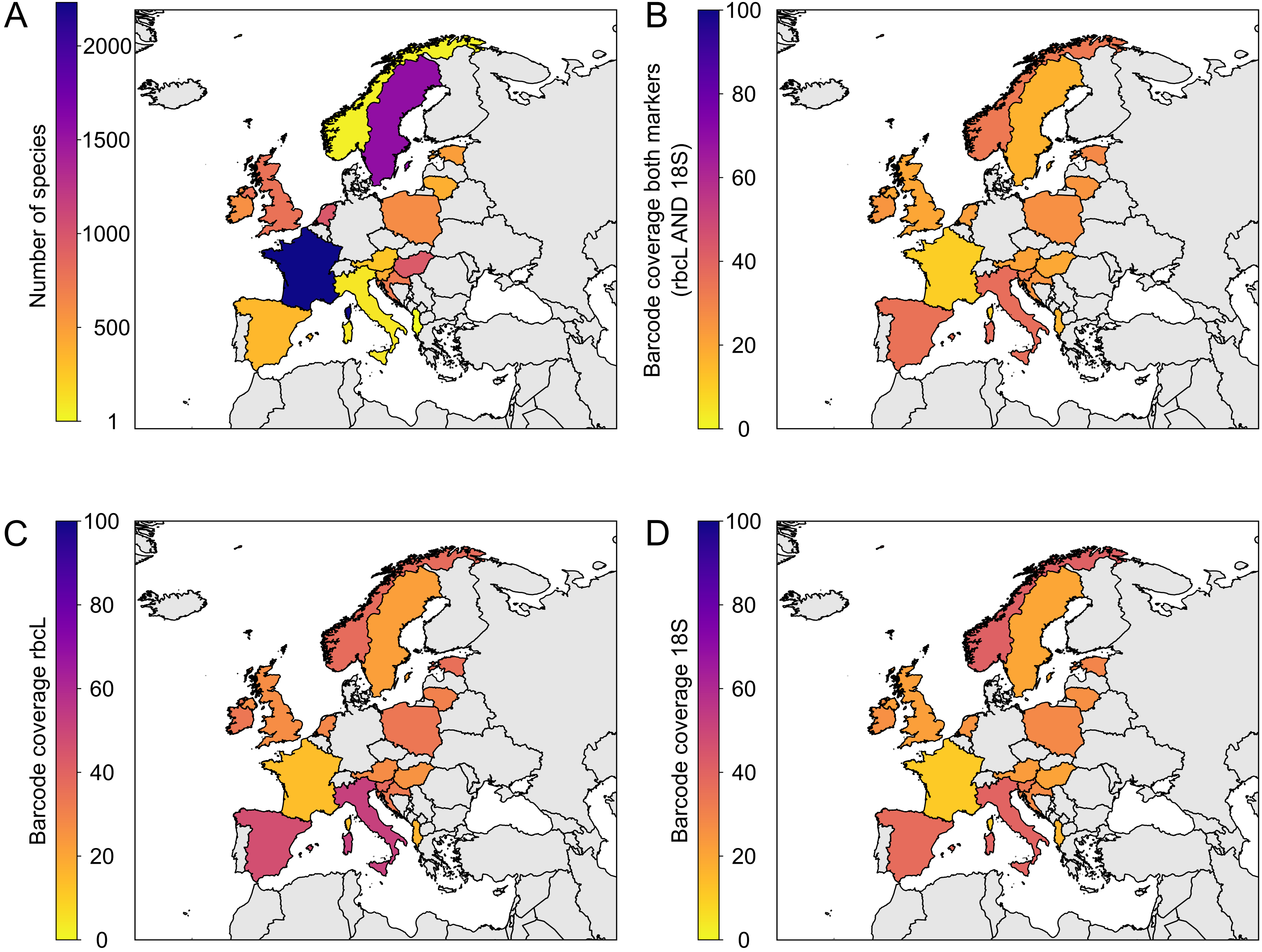

### Supplement Fig. 2

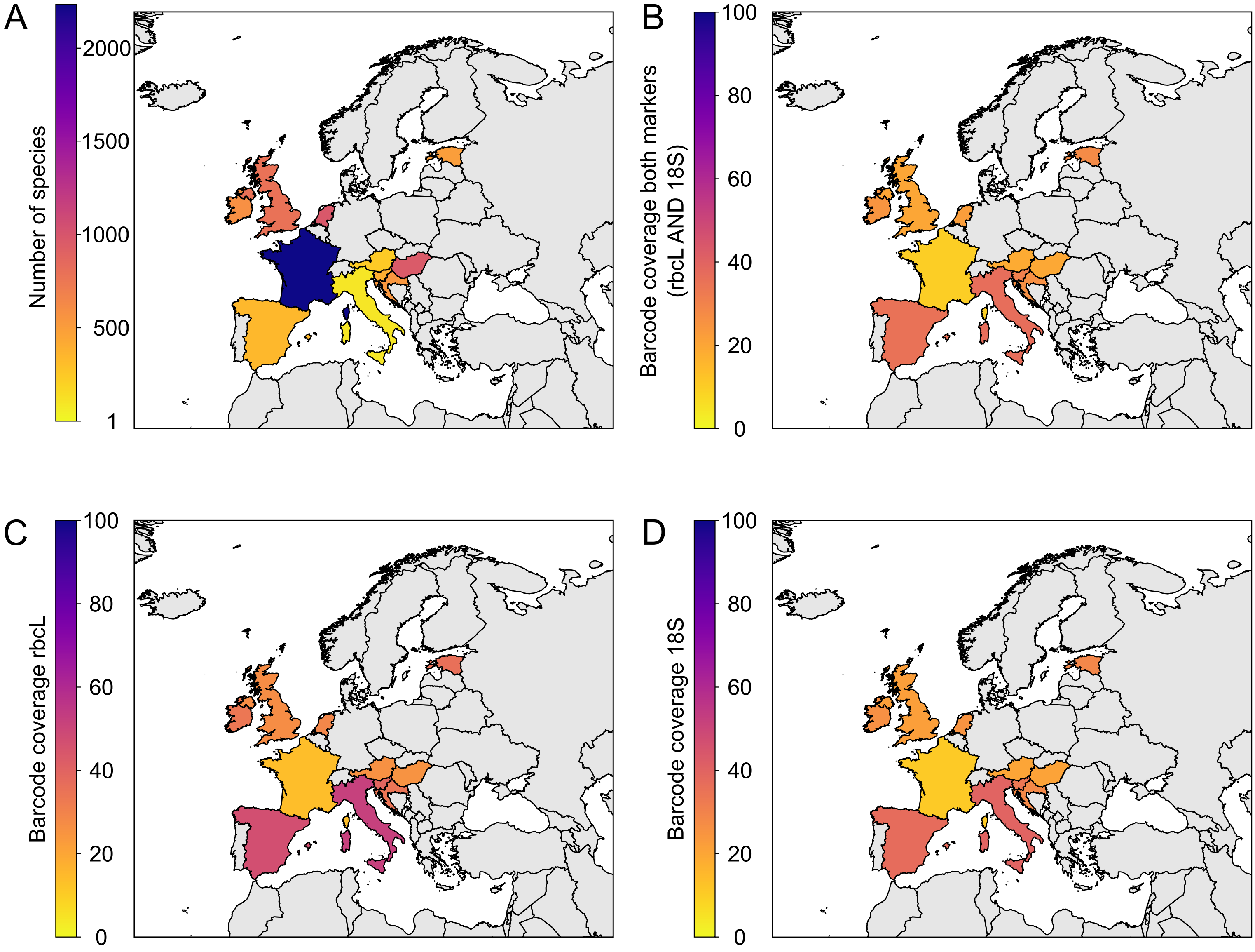

### Supplement Fig. 3

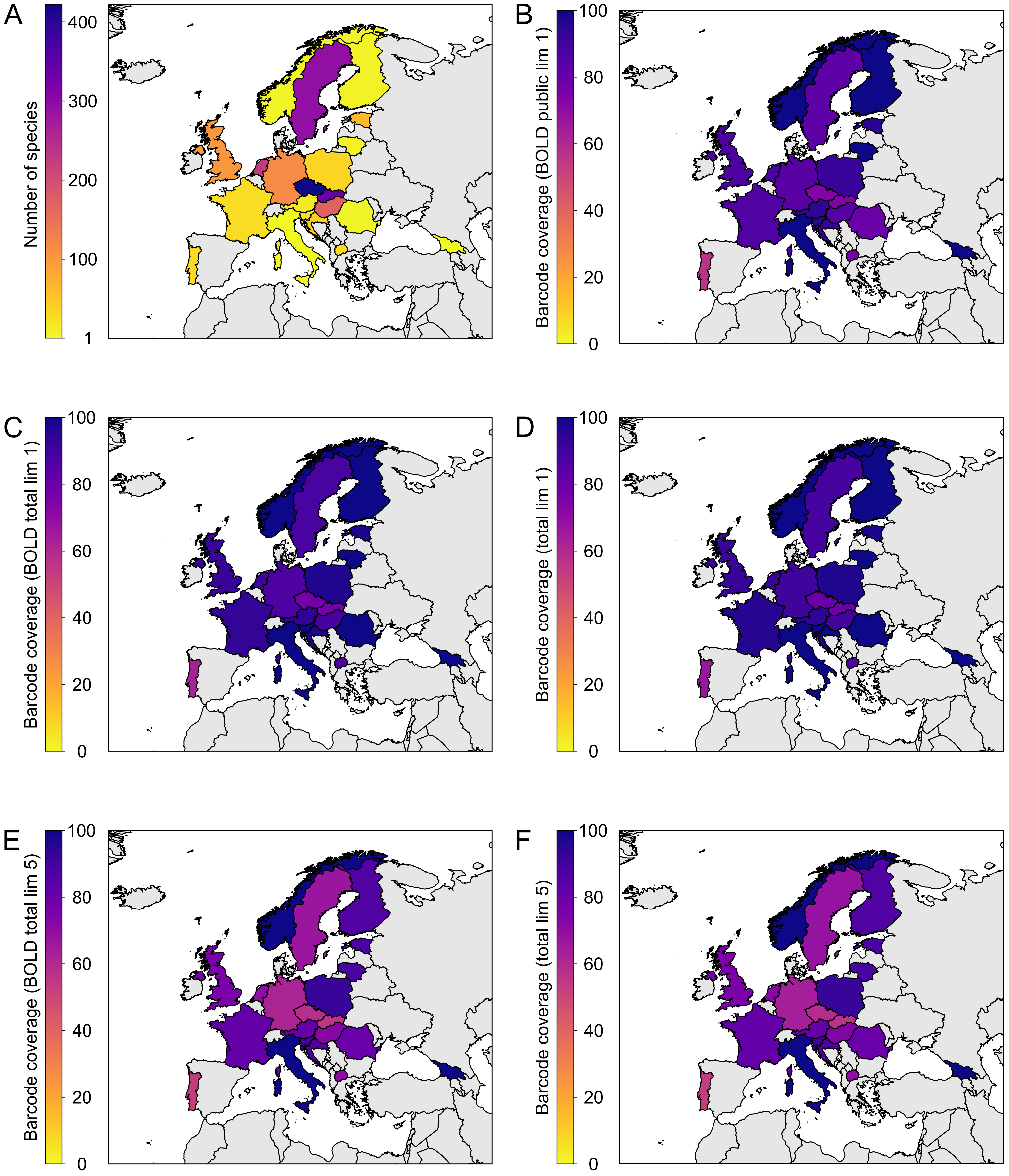

### Supplement Fig. 4

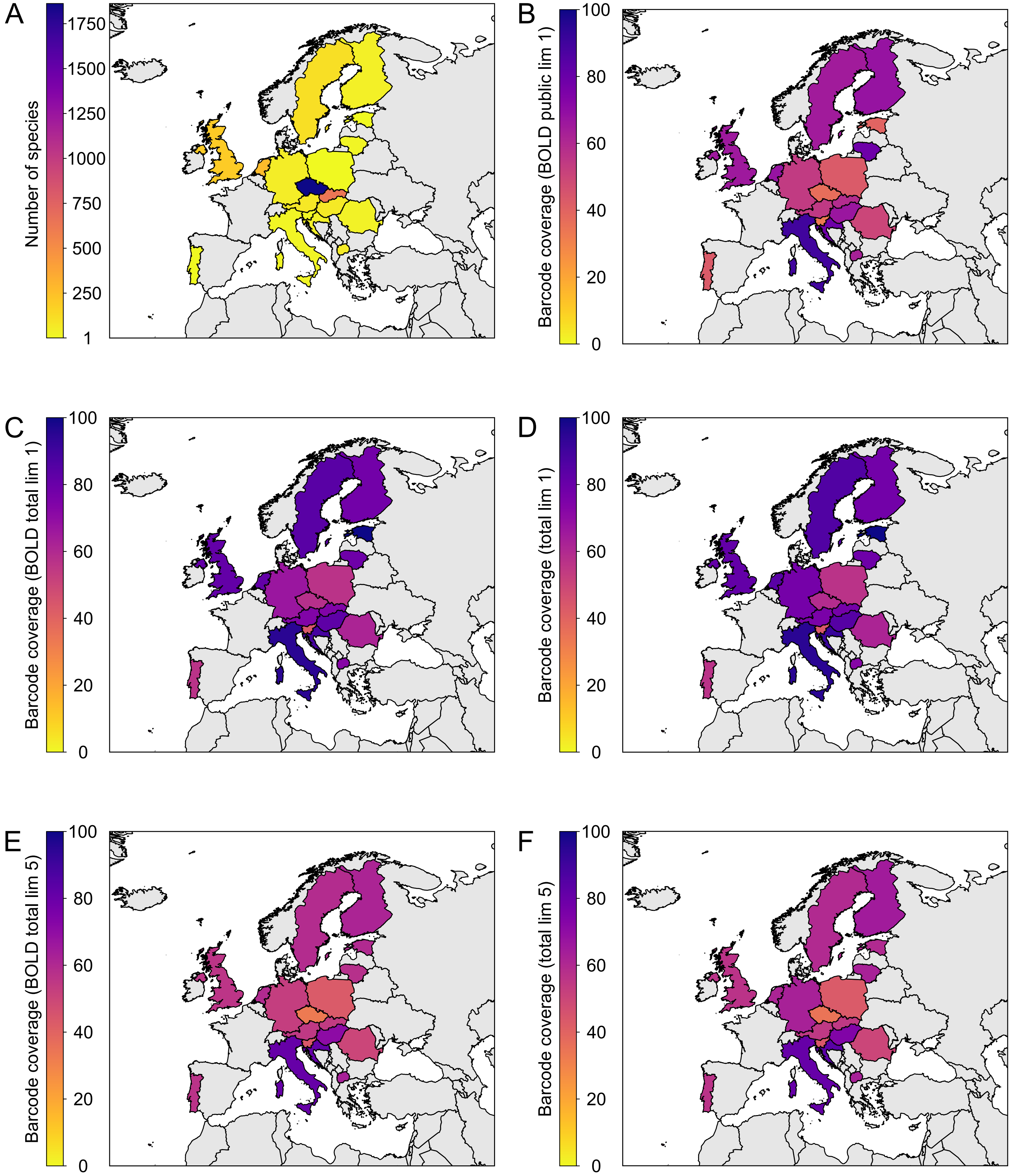

### Supplement Fig. 5

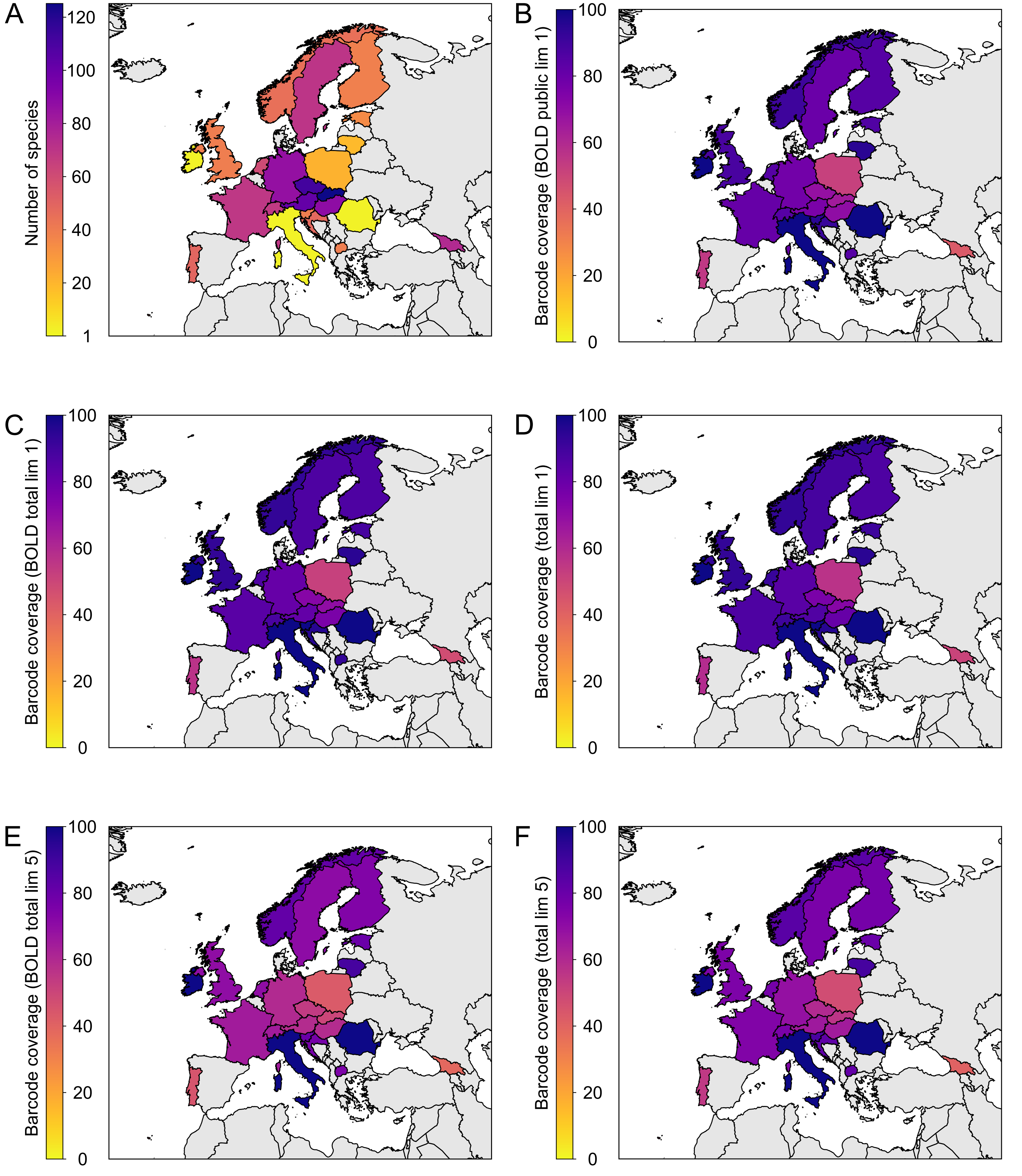

### Supplement Fig. 6

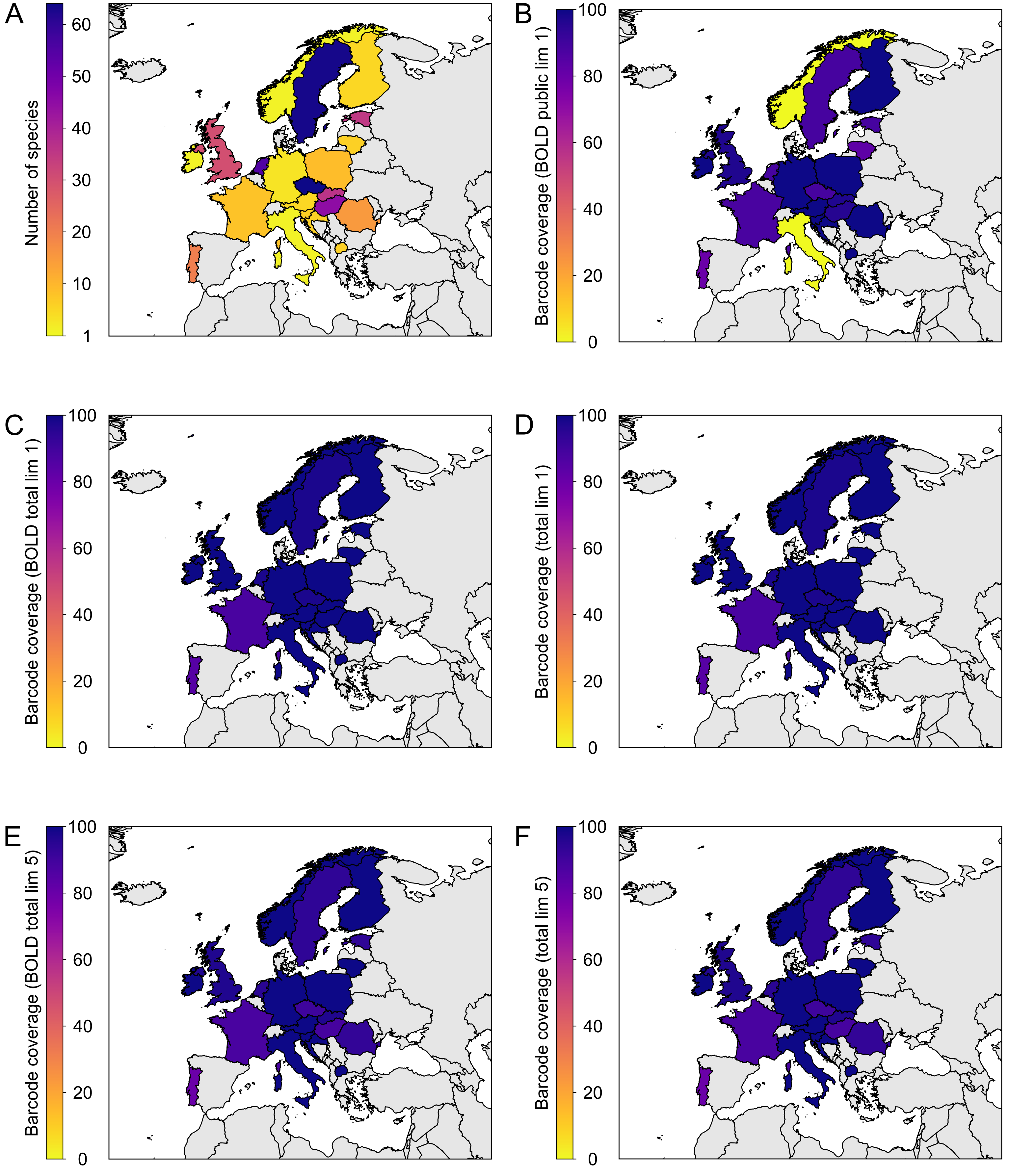

### Supplement Fig. 7

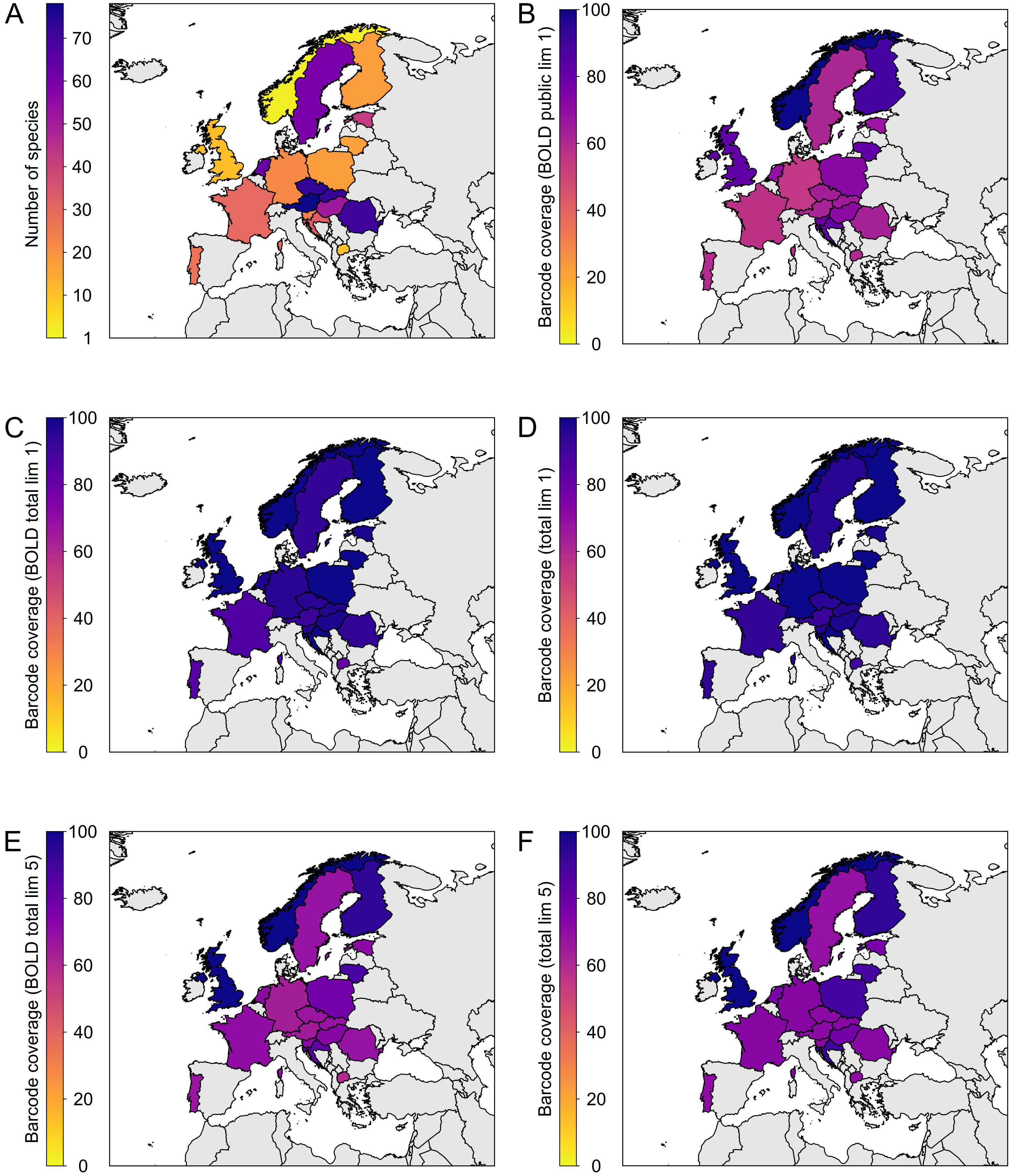

### Supplement Fig. 8

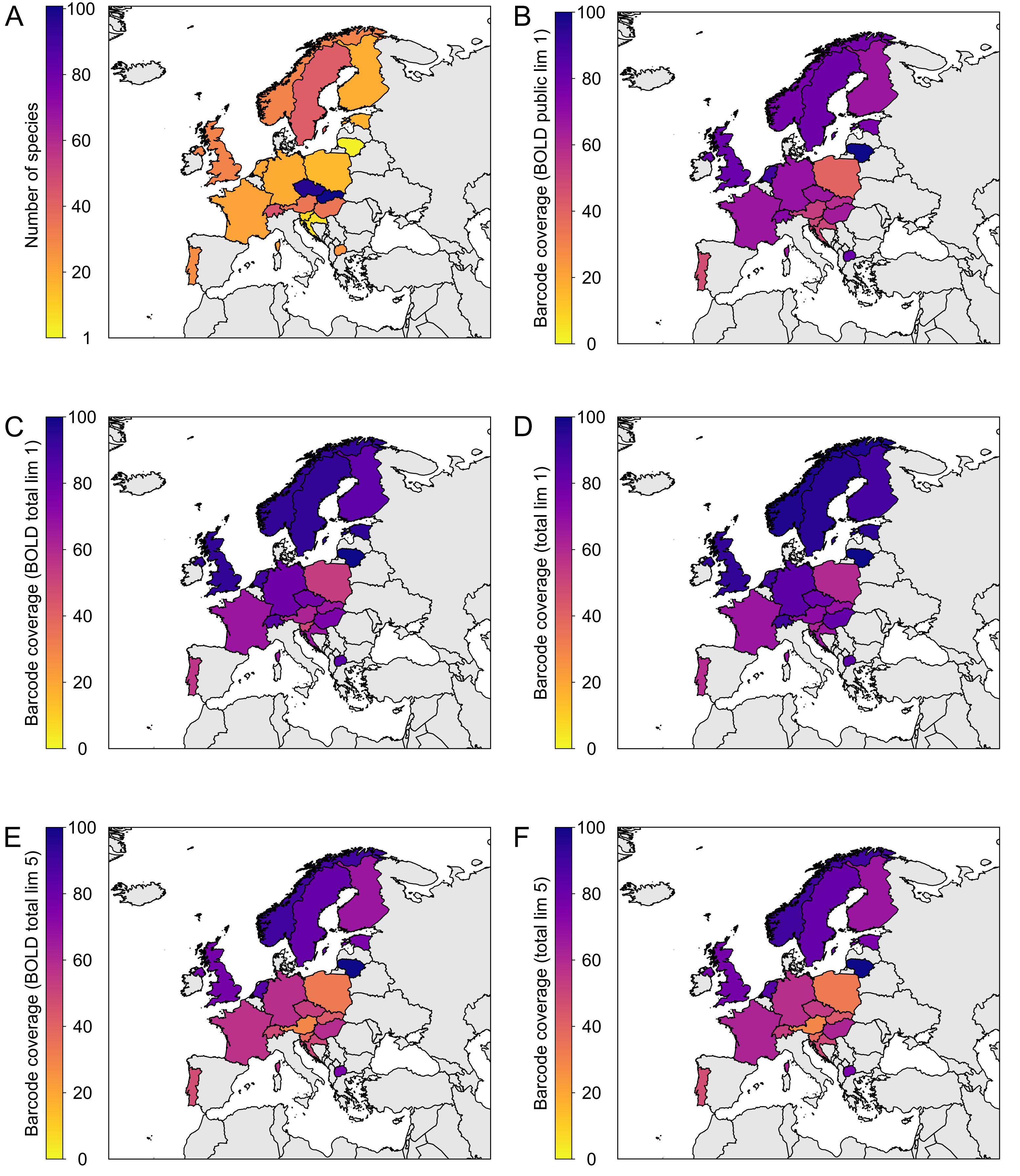

### Supplement Fig. 9

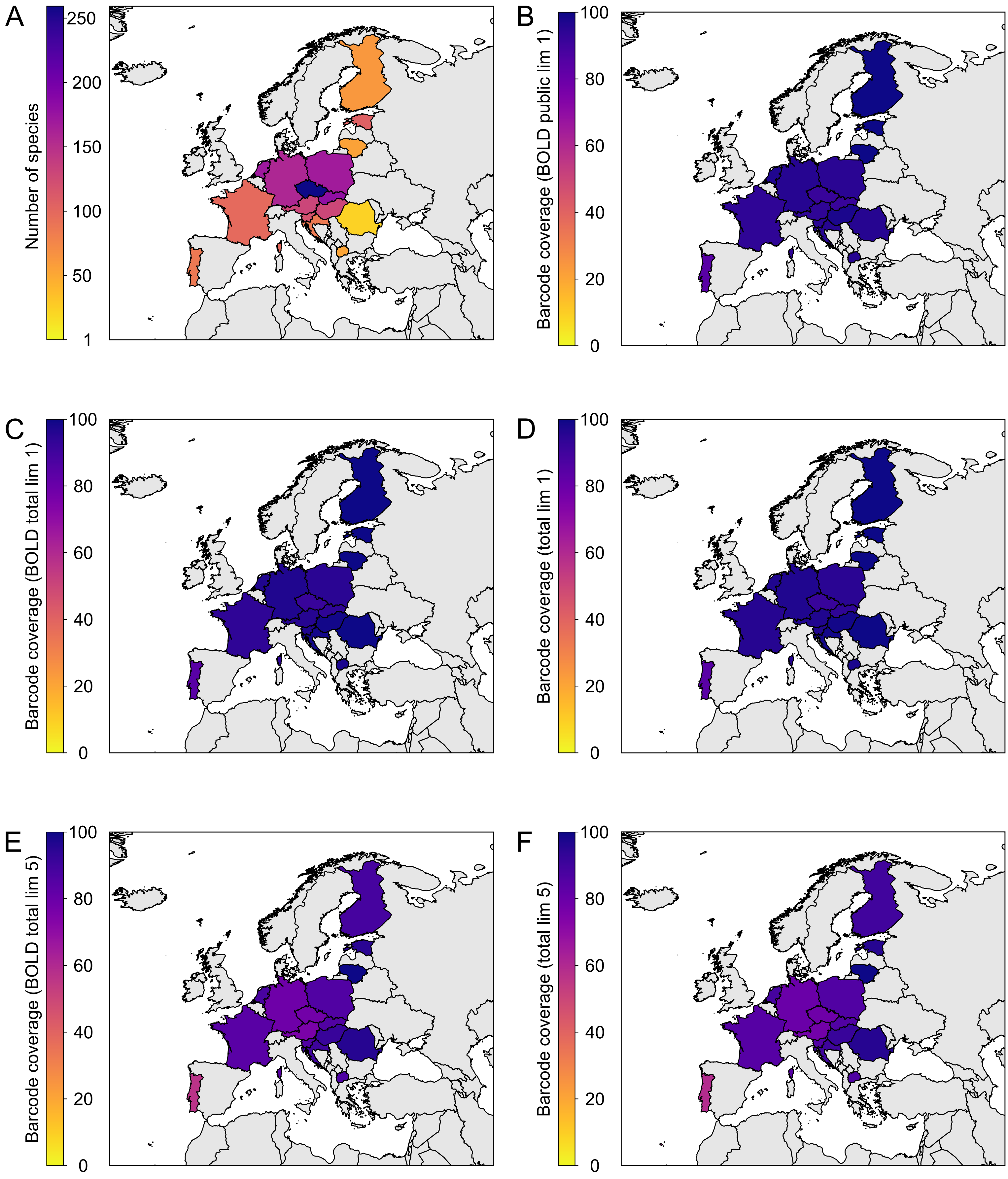
